# SBP transcription factors repress maize brace root initiation and regulate primordia expansion

**DOI:** 10.64898/2026.11.17.767364

**Authors:** Thanduanlung Kamei, Ashley N. Hostetler, Sarah Tallents, Taran A. Kermani, Ed Bridge, Annis E. Richardson, Norman B. Best, Erin E. Sparks

**Affiliations:** Department of Plant and Soil Sciences, University of Delaware, Newark, DE, USA; Division of Plant Science and Technology, University of Missouri, Columbia, MO, USA; Institute of Molecular Plant Science, School of Biological Sciences, University of Edinburgh, Edinburgh, UK; USDA-ARS, Plant Genetics Research Unit, Columbia, MO, USA; Bond Life Sciences Center, University of Missouri, Columbia, MO, USA; Donald Danforth Plant Science Center, St. Louis, MO, USA

## Abstract

Nodal roots are critical determinants of maize productivity and resilience, but the molecular mechanisms governing their development remain poorly defined. Transcriptome analyses across three stages of maize brace root (aboveground nodal roots) development identified the squamosa promoter binding protein (SBP) transcription factors *tasselsheath4* (TSH4), *unbranched2* (UB2), and *unbranched3* (UB3) as negative regulators of brace root initiation. Analysis of single, double, and triple mutants revealed a dose-dependent increase in the number of nodes with brace roots and a non-linear increase in the number of brace roots per node. The increased number of brace roots within a node resulted from a physical expansion of the stem pulvinus and de-repression of brace root initiation programs, which resulted in two tiers of brace roots at each node. Downstream analyses revealed that SBP TFs negatively regulate *liguleless2 e*xpression, which acts in concert with *liguleless1* to promote brace root initiation, and phytohormone analysis identified a mechanism whereby *sbp* mutants exhibited increased ACC accumulation. Differential levels of gibberellin (GA_4_) were identified between *tsh4* and *ub2;ub3* mutant nodes. The differential levels of gibberellin regulated primordia expansion resulting in distinct brace root phenotypes – *tsh4* mutants had fewer, thicker brace roots whereas *ub3* mutants had more and thinner brace roots. The *tsh4* and *ub3* GA_4_-mediated post-initiation expansion mechanism was also found to underlie natural variation in brace root size and number. Together, our findings identify SBP transcription factors as central regulators of brace root initiation and primordia expansion and show that altering developmental trajectories can alter brace root architecture.

## Introduction

Root systems regulate water uptake, nutrient acquisition, microbial interactions, and plant anchorage. In maize, the root system is composed of seed-borne, stem-borne, and root-borne roots. Seed-borne roots include the primary and seminal roots, stem-borne roots are also called nodal roots and include crown and brace roots, and root-borne roots are lateral roots that develop from all other root types. Nodal roots are the predominant functional root type after the vegetative leaf 6 stage (V6) (Hochholdinger et al. 2004). Nodal roots develop sequentially along the stem starting with crown roots originating from belowground nodes and continuing with brace roots originating from aboveground nodes (Blizard and Sparks 2018). Modulating the number and size of nodal roots is an important factor for survival under nutrient limited conditions (Schneider et al. 2021) and promoting plant anchorage, preventing root lodging (Hostetler et al. 2022). However, targeted improvement of root systems requires a detailed understanding of the developmental pathways that modulate the number and size of nodal roots.

Developmentally, nodal roots arise from the stem at the base of an internode directly above the node (Brown et al. 1959; Lamb et al. 2024). For brace roots, this region is the stem pulvinus, a tissue traditionally associated with gravitropic bending responses (Perera et al. 1999). Because the maize stem pulvinus has been characterized primarily based on its gravitropic response, it remains unclear whether a homologous structure exists in the belowground stem where crown roots initiate and internodes are highly compressed. Despite these potential differences, the developmental mechanisms that regulate nodal root development are thought to be shared between crown and brace roots. The four sequential stages of nodal root development have been defined as 1) induction, 2) initiation, 3) emergence, and 4) growth (Bellini et al. 2014; Hostetler et al. 2021; Kamei et al. 2024). During induction, nodal root founder cells are defined by their competence to undergo cell division. It is currently unknown when and where nodal root founder cells arise. Initiation then occurs when the founder cells divide to give rise to the nodal root primordia. Emergence occurs when nodal root primordia expand and emerge from the stem. After emergence, the growth stage determines the rate and direction of subsequent root growth.

Each stage of nodal root development is governed by distinct transcriptional and phytohormone networks, although the molecular mechanisms regulating each stage remain poorly defined. To date, fewer than 25 maize mutants have been reported with altered nodal root phenotypes. Many of these mutants alter the number of nodes that produce roots, with the genes underlying these mutants often controlling the juvenile-to-adult phase transition (Hostetler et al. 2021). Mutants with delayed phase transitions have more nodes with roots, whereas mutants with accelerated phase change transitions have less nodes with roots (Hostetler et al. 2021). Among these mutants is *Corngrass1 (Cg1)*, which remains in the juvenile phase and develops roots at all nodes (Chuck et al. 2007). *Cg1* is a dominant mutant caused by the overexpression of microRNA156 (miR156), which targets the degradation of squamosa promoter binding protein (SBP) transcription factors (TFs) (Wei et al. 2018). Among the SBP TFs that are targeted by miR156, an RNAi knock-down of *teosinte glume architecture 1* (*tga1)* increased brace root node number (Wang et al. 2015). This SBP TF is a key domestication gene that resulted in the loss of seed cases (Wang et al. 2015). A separate group of SBP TFs, *tasselsheath4* (TSH4), *unbranched2* (UB2) and UB3, have been identified as regulators of inflorescence initiation and branch number (Chuck et al. 2010, 2014; Du et al. 2020; Dong et al. 2024). These three SBP TFs are closely related, with UB2 and UB3 having high sequence similarity and considered duplicated in maize (Chuck et al. 2014). Mutants of *tsh4* have reduced tassel branch number and increased bract outgrowth (Chuck et al. 2010). Bracts are modified leaves located at the base of inflorescence branches whose development is normally suppressed in maize. Single mutants of either *ub2* or *ub3* have a mild reduction in tassel branch number, while the double mutant (*ub2;ub3*) shows a more severe reduction like *tsh4* (Chuck et al. 2014). In contrast to *tsh4,* the *ub2;ub3* mutants do not show bract de-repression but have reduced kernel row number (Chuck et al. 2014). In addition to the changes observed in lower order mutants, the *tsh4;ub2;ub3* triple mutant increased tiller number, highlighting the combined role of these SBP TFs in the repression of tiller bud outgrowth (Chuck et al. 2014). These results emphasize the nuances of SBP TFs functioning both independently and together to coordinate meristem activity and branch number. The function of SBP TFs in nodal root meristem activity and branching have not been previously reported and are the basis of this study.

Other mutants that alter nodal root development implicate phytohormone pathways. For example, the auxin-responsive *rootless concerning crown and seminal roots* (RTCS) TF is a critical regulator of early root development, with the *rtcs* mutant having no nodal or seminal root primordia (Taramino et al. 2007). This mutant indicates that auxin positively regulates early nodal root induction or initiation. However, another auxin mutant, *rootless with undetectable meristem1* (*rum1)* which encodes an AUX/IAA auxin signaling protein, has reduced seminal and lateral root development, but does not impact nodal roots (von Behrens et al. 2011; Zhang et al. 2015), demonstrating the specificity of auxin signals varies by root development program. Other studies have implicated ethylene as a positive regulator of nodal root emergence. Specifically, the overexpression of *auxin-regulated gene involved in organ size8* (*argos8*) inhibits ethylene signaling and reduces nodal root emergence (Shi et al. 2019). Two other mutants, *rootless1* and *rap2.7*, disrupt genes encoding ethylene-responsive TFs and reduce the number of nodes with roots, although the stage(s) of development that are disrupted have not been reported (Jenkins 1930; Sakai and Taramino 2009; Li et al. 2019). A recent study looked at phytohormone levels in aboveground nodes of maize plants grown under varying soil compaction and water content conditions to investigate the hormonal regulation of brace root development (Wei et al. 2026). In this study, ethylene was positively associated with brace root development, cytokinin was negatively associated with brace root development, and auxin had minimal effects (Wei et al. 2026). Thus, while phytohormones likely play an important role in nodal root development, there has been a limited quantification of these interactions.

In this study, we defined the regulation of nodal root development using three stages of brace root development (induction, initiation, and emergence). Transcriptome profiling and differential gene analyses identified the SBP TFs TSH4, UB2, and UB3 as candidate negative regulators of brace root initiation. Analysis of mutants for these SBP TFs revealed two distinct brace root phenotypes – more nodes that produced brace roots and an extra tier of brace roots at each node. The extra tier of brace roots was the result of an expanded competence zone (i.e., stem pulvinus) and de-repression of brace root initiation programs. Surprisingly, this phenotype was not found in crown roots, refuting the idea that nodal root developmental programs are generalizable. We found that TSH4 and UB2/UB3 repress ethylene and cell cycle genes to limit brace root initiation. Phytohormone profiling also identified differences in gibberellin (GA_4_) levels between *tsh4* and *ub2;ub3* mutants. Differences in GA_4_ levels resulted in two distinct strategies of node occupancy – *tsh4* having fewer and thicker brace roots and *ub3* having more and thinner brace roots. Variation in the relative expression of *tsh4* and *ub3* was found to underlie natural variation in brace root number and size. Collectively, these results form the foundation for modulating development to optimize nodal root function in maize.

## MATERIAL AND METHODS

### Plant growth conditions

Maize inbred (B73, W22, CML333, and PHHB4) and mutant (*tsh4-mum1* in W22*, ub3-mum3* in W22*, ub2-mum1;ub3-mum3* in W22*, ub2-mum1;ub3-mum3;tsh4-mum1* in B73, *ub2-mum1;ub3-mum3;tsh4-mum1* in W22 (Chuck et al. 2010, 2014), *liguleless1* (*lg1*), *lg2,* and *lg1;lg2* in B73 (Becraft et al. 1990; Harper and Freeling 1996), *dwarf3* (*d3*) in B73 (Winkler and Helentjaris 1995)), lines were used for this study.

Greenhouse experiments were conducted for assessment of histology, gene expression profiling, *Cg1* and *lg2* assessment, and phytohormone profiling. For all greenhouse experiments except *lg2* phenotyping, plants were grown in 11.4-L air pruning pots (Superoots Air-Pots) filled with PRO-MIX BK55 soil (histology, gene expression profiling, and *Cg1* assessment) or PRO-MIX BX soil (phytohormone profiling). Plants were supplied with Peters Excel 15-5-15 nutrient solution at 200 ppm nitrogen. Supplemental lighting was provided with overhead high-pressure sodium lamps was set at 50 KLUX (∼ 925.93 µmol/m^-2^/s^-1^ PPFD), for a 16h photoperiod and with daytime and nighttime temperatures maintained at 26.7°C (80°F) and 21.1°C (70°F), respectively. For *lg2* phenotyping, plants were grown in 5-L pots filled with peat-free soil plus osmocote, for a 16h photoperiod and with daytime and nighttime temperatures maintained at 28°C (82.4°F) and 23°C (73.4°F), respectively. Supplemental lighting was provided by overhead Valoya RX400 NS12 lights.

Field experiments were conducted in Newark, DE, USA (2023 and 2024) and Columbia, MO, USA (2026). Seeds were planted on 17-May-2023, 21-May-2024, 14-May-2026 at a density of 18 plants per 3.66-meter single-row plot with 0.762-meter row spacing (6.4 plants m⁻²). For DE, pre-emergence herbicide (Lexar at 8.18 L ha^-1^ and Simazine at 2.81 L ha^-1^), post-emergence herbicide (Accent at 0.05 L ha^-1^), soil insecticide (COUNTER 20 G at 6.16 kg ha^-1^), and fertilizer (ammonium sulfate 21-0-0 at 100.88 kg ha^-1^) were applied at planting. Fields were side-dressed with 30% urea ammonium nitrate (at 374.16 L ha^-1^) approximately one month after planting. Weather data can be found at the Delaware Environmental Observing System (https://www.deos.udel.edu/) by selecting the Newark, DE-Ag Farm station. For MO, pre-emergence herbicide (Degree Xtra at 8.65 L ha^-1^), post-emergence herbicide (Atrazine at 2.34 L ha^-1^), and fertilizer (NPK 160-0-40 at 179.3 kg N ha^-1^ and 44.8 kg K ha^-1^ pre-planting) was applied. Weather data can be found at Weather Underground (https://www.wunderground.com/dashboard/pws/KMOCOLUM110). In 2022, *lg1*, *lg2*, and the double *lg1;lg2* mutants were grown in the field at the University of Massachusetts Amherst Crop and Animal Research and Education Farm in South Deerfield, MA, USA.

For assessing crown root phenotypes in seedlings, W22, B73, *tsh4, ub3, ub2;ub3, and ub2;ub3;tsh4* seeds were surface sterilized with a 50% (v/v) sodium hypochlorite (bleach) solution for 15 minutes. Seeds were rinsed with water five times and placed on sterile germination paper (Anchor paper Co 38 Reg Germ paper 12x24 2000EA). The paper was rolled and placed in 1 L beakers containing 200 mL of sterile distilled water. Beakers were transferred to a tabletop growth chamber (Seed and herb domed propagator 9”D x 16”W x 14.5”H).

### Histology

The three lowest aboveground nodes (closest to the soil) of B73 plants were isolated at the vegetative leaf 4 (V4), V6, V8, and V10 stage. Nodes were hand sectioned with a razor blade and fixed overnight at 4℃ in 4% paraformaldehyde (PFA) in 1X phosphate-buffered saline (PBS). Sections were rinsed and stained with 0.2% fuchsin in Clearsee (Ursache et al. 2018) and imaged on a Zeiss Axiozoom microscope with 2X magnification. The number of primordia were counted for each node, and the length of primordia was measured from base to tip using ImageJ (Schindelin et al. 2015; Rueden et al. 2017).

### Transcriptomics

The three aboveground nodes closest to the soil of B73 plants at the V6 stage were dissected, flash-frozen, and processed for total RNA extraction. The second lowest aboveground node of *tsh4* and *ub2;ub3* mutant plants at the V6 stage were dissected into lower tier (T1) and upper tier (T2) regions. For W22 plants at the V6 stage, the same node was dissected, but only from tissue directly above the pulvinus (T1). Total RNA was extracted using the Qiagen RNeasy Mini Kit and mRNA was purified using the NEBNext® Poly(A) mRNA Magnetic Isolation Module according to the manufacturer’s protocol. Strand-specific mRNA libraries were generated using the YourSeq (FT & 3’DGE) RNAseq Library Kit (Amaryllis Nucleics) following the manufacturer’s instructions. B73 libraries were pooled and sequenced on an Illumina NextSeq 500 (High Output v2 75-cycle kit) to generate 80-bp single-end reads through Amaryllis Nucleics. Mutant and W22 libraries were pooled and sequenced on an Element AVITI (150-cycle kit) to generate 150-bp single-end reads at the University of Delaware Core DNA Sequencing Facility).

Quality and mapping analyses were completed in Galaxy (Galaxy Community 2024). In brief, quality checks were assessed by FastQC (version 0.12.1) (Wingett and Andrews 2018) and MultiQC (1.27+galaxy1) (Ewels et al. 2016) with all default settings. The adapter and PolyA tail were trimmed using Cutadapt (4.9+galaxy1) (Martin 2011). Reads were mapped to the *Zea mays* Zm-B73-REFERENCE-NAM-5.0 reference sequence (for B73 samples) and Zm-W22-REFERENCE-NRGENE-2.0 (for W22 and mutant samples) using HISAT2 (2.2.1+galaxy1) (Kim et al. 2015). Read counts were generated using featureCounts (2.0.8+galaxy0) (Liao et al. 2014). The R package DESeq2 (version 1.44.0)(Love et al. 2014) was used to identify differentially expressed genes using a negative binomial generalized linear model. Differentially expressed genes were filtered using a minimum normalized counts > 1 in all samples, a false discovery rate (FDR) cutoff < 0.01, and a log2 fold change > |1|. Annotated transcriptional regulators were downloaded from Grassius (Yilmaz et al. 2009), and the maize genome version 5 classification was used to assign TF gene families. Gene identifiers, gene symbols, and gene annotations were obtained from MaizeGDB (Woodhouse et al. 2025). Principal component analysis (PCA) was performed using the *stats* package in R version 4.4.0 (2024). Heatmaps were generated in the *pheatmap* package version 1.0.13 (Kolde 2025).

### Phenotyping

Brace root node number, brace root number, and pulvinus length were quantified in 2024 for W22, *tsh4, ub3, ub2;ub3, ub2;ub3;tsh4* plants. At 71 days after planting (dap), the total number of aboveground nodes containing brace roots (brace root primordia and emerged brace roots) was manually recorded for 8-10 plants per genotype. At 48, 58, and 65 dap, images of the base of plants (n = 2-24) were captured. From images, the total number of brace roots at aboveground nodes 3 and 4, and the total number of brace roots at Tier 1 (T1) and Tier 2 (T1) (aboveground nodes 3 and 4) were quantified. Due to differences in developmental timing, W22 node 3 and node 4 were quantified at 65 dap and mutants were quantified at 48 dap for node 3 and 58 dap for node 4. Pulvinus length of aboveground node 4 was quantified at 48 dap. Image analysis was conducted using Fiji (version 1.54p) (Schindelin et al. 2015). Two measurements were taken per plant and averaged.

Brace root size and number were quantified in 2023 for *tsh4* and *ub2;ub3* (n=7) at 119 dap. Brace root diameter was averaged from four brace roots excised from aboveground node 2 and measured with calipers. The number of brace roots per node were manually recorded for aboveground nodes 2 and 3. Brace root size and number, and stem diameter were quantified in 2026 from images of *d3* mutant plants (n=3) and wild-type siblings (n=12) at 61 dap. From the images, the total number of brace roots at aboveground node 2 were quantified. The width of the front-most brace root and stem width were quantified using Fiji. The total number of aboveground nodes containing brace roots (brace root primordia and emerged brace roots) was manually recorded for *ub2;ub3;tsh4* in B73 (n=4), B73 (n=12), *ub2;ub3;tsh4* in W22 (n=7), and W22 (n=19) at 69 dap in 2023.

Brace root tier number was quantified from images of greenhouse-grown *Cg1* mutants and wild-type siblings at 34 dap in 2024. Brace root node number was assessed from images of field-grown *lg1, lg2, lg1;lg2*, and wild-type siblings (n = 2–7) at 98 dap in 2022. Emerged brace roots were counted at aboveground nodes 1–4 in greenhouse-grown *lg2* mutants (n = 6) and wild-type siblings (n = 3) at 115 dap in 2026.

### TSH4 and UB3 downstream targets

TSH4 and UB3 downstream targets were evaluated as shared with ear DNA binding targets and unique from ear DNA binding targets. For shared targets, previously published UB3 DNA Affinity Purification sequencing (DAP-seq; (Ricci et al. 2019; Dong et al. 2024) and TSH4 Chromatin Immunoprecipitation sequencing (ChIP-seq; (Dong et al. 2024)) datasets from immature ears were integrated with brace root transcriptome datasets. UB3 DAP-seq targets and TSH4 ChIP-seq targets were first intersected to identify genes bound by both TFs in immature ears. This shared target list was then intersected with brace root initiation candidates (primordia/no primordia). To determine if these shared targets were also regulated during brace root initiation, RNA-seq datasets from *tsh4* and *ub2;ub3* plants were analyzed. Specifically, differential gene expression analyses were performed where T1 and T2 of each mutant was compared to W22 T1. Genes that were differentially expressed in all four comparisons (*tsh4* T1/W22 T1; *tsh4* T2/W22 T1; *ub2;ub3* T1/W22 T1; *ub2;ub3* T2/W22 T1) were identified by intersection and subsequently compared with the SBP TFs DNA binding targets in immature ears.

To identify downstream targets that were unique to brace root initiation, the DEG lists for the four mutant comparisons with W22 (*tsh4* T1/W22 T1; *tsh4* T2/W22 T1; *ub2;ub3* T1/W22 T1; *ub2;ub3* T2/W22 T1) were again intersected and compared with candidate regulators of brace root initiation (primordia/no primordia). Those that were shared among the five RNA-seq datasets were considered candidates for downstream targets of SBP TFs during brace root initiation.

### Phytohormone analysis

Tier 1 tissue from the second aboveground node of W22, *tsh4* and *ub2;ub3* was dissected as described for transcriptome analysis. Dissected samples were ground into fine powder in liquid nitrogen. Approximately 100 mg of ground tissue was subjected to LC/MS/MS analysis as described in (Dilkes and Best 2026).

### RT-qPCR

To identify genotypes varying for brace root traits (diameter and number) but not stem diameter, previously published phenotypic data from 52 maize inbred lines was assessed (Hostetler et al. 2022). From this analysis, two genotypes (CML333 and PHHB4) with statistically similar stem diameters but contrasting brace root strategies (fewer and thicker brace roots versus more and thinner brace roots) were selected for further analysis.

Quantitative PCR (qPCR) was performed to calculate the relative expression of *tsh4* and *ub3* in the aboveground node 2 of CML333 and PHHB4. RNA was isolated using the Zymo Quick-RNA™ Plant Miniprep. First stand cDNA was synthesized using the SuperScript™ VILO™ cDNA Synthesis Kit. Primers were designed for *tsh4* and *ub3* using the Primer3Plus online software (Untergasser et al. 2012)(**Table S1**). qPCR reactions were performed using Thermo Fisher Scientific SYBR Green Universal Master Mix on a Thermo Fisher Scientific QuantStudio3 real-time PCR system with three technical and three biological replicates. Relative gene expression levels were calculated using the ddCT method (Livak and Schmittgen 2001).

### Statistics

All statistical analyses were performed in R version 4.4.0 (R Core Team 2024) and all figures were generated with the *ggplot* version 4.0.3 (Wickham 2016) and *cowplot* version 1.2.0 (Wilke 2015) packages. To assess differences in traits, analysis of variance (ANOVA) models were used. Prior to running each ANOVA, the normality of residuals was assessed with a Shapiro–Wilk test from the *stats* package. If residuals were not drawn from a normal distribution (primordia size and stem width), a Tukey’s Ladder of Powers transformation (*rcompanion* package version 2.4.1)(Mangiafico 2021) was used, and the transformed data was used in the ANOVA. When *P*<0.05, pairwise means were compared using the *agricolae* package version 1.3.7 (de Mendiburu 2023). To assess genotypic (independent variable) differences in traits, individual one-way ANOVA models were run with the following dependent variables: brace root node count; total brace root count in *sbp* mutants and W22 (aboveground nodes 3 and 4, analyzed separately); tier 1 brace root count (aboveground nodes 3 and 4, analyzed separately); tier 2 brace root count (aboveground nodes 3 and 4, analyzed separately); pulvinus length; phytohormone abundance (each hormone analyzed separately); brace root diameter; and stem diameter. For brace root node count, all three datasets were analyzed separately: *sbp* mutants in W22 background; *liguleless* mutants (*lg1, lg2, and lg1;lg2*) in B73; and the triple mutant (*ub2;ub3;tsh4*) in B73 and W22. For brace root diameter, the *sbp* mutants (*tsh4 and ub2;ub3*) and the *d3* mutant were analyzed separately. For stem diameter, the *d3* mutant and 52 maize inbred lines (Hostetler et al. 2022) were analyzed separately. To assess differences in brace root number between *lg2* and B73, the following two-way ANOVA was run: brace root count ∼ genotype * node of origin. To assess developmental differences in brace root development, the following two-way ANOVA was run: brace root primordia count ∼ vegetative leaf stage * node of origin.

## RESULTS

### SBP transcription factors are candidate regulators of brace root initiation

The spatiotemporal progression of maize brace root development was quantified at the vegetative leaf 4 (V4) to V10 growth stages for the three aboveground nodes closest to the soil (**Fig S1**). Brace root development could be observed across two temporal trajectories: within individual nodes over time and sequentially across nodes within a growth stage (**Fig S1**). Among the growth stages, the V6 stage captured three stages of brace root development: 1) induction stage, the top node with no primordia; 2) initiation stage, the middle node with primordia; 3) emergence stage, the lowest node with brace roots emerging (**Fig 1A**).

**Fig 1.**
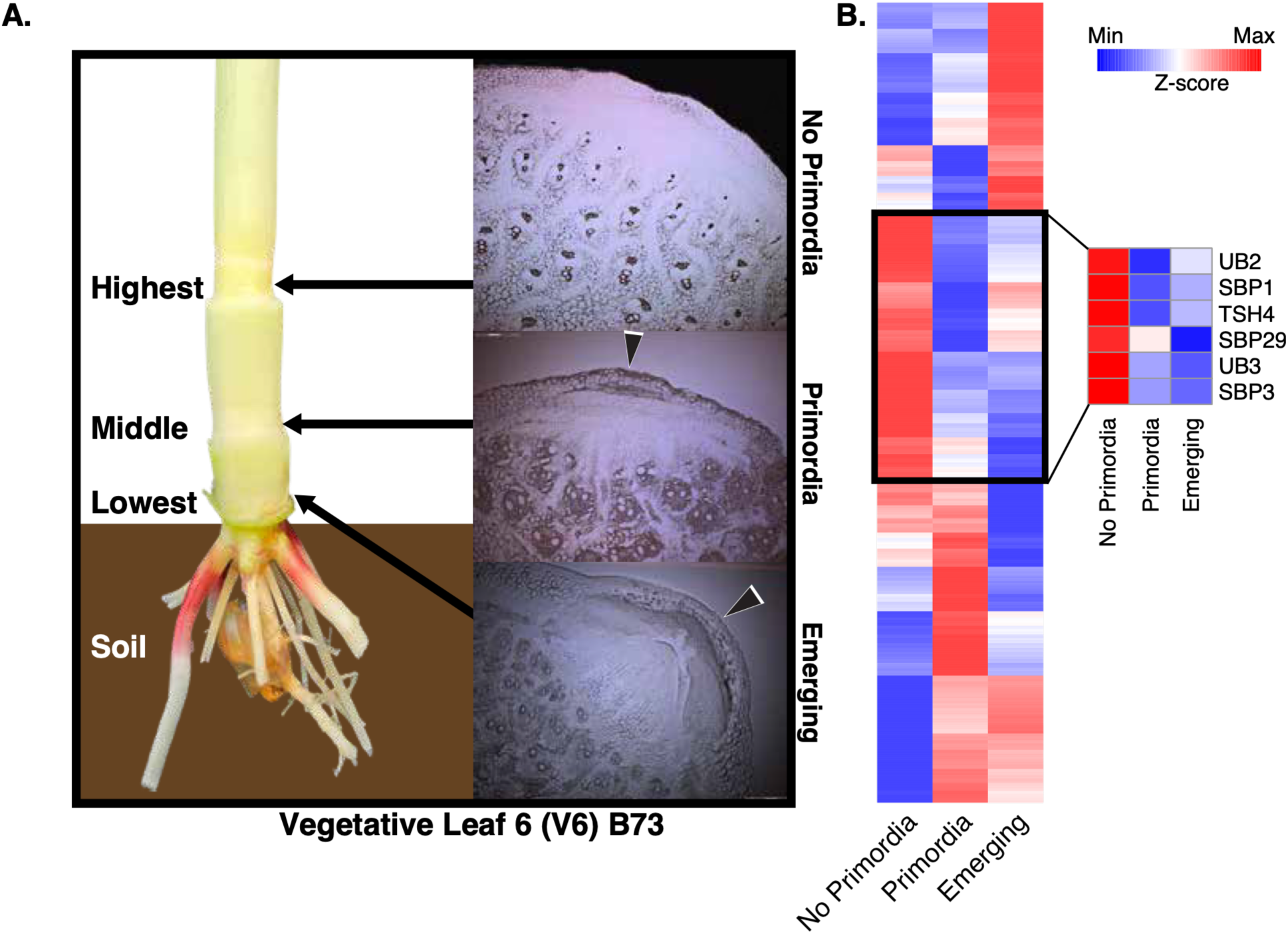
Node-specific transcriptome signatures define brace root developmental stages. A) A B73 V6 plant demonstrating the three stages of brace root development (no primordia, primordia, and emerging) that are observed at the lowest three aboveground nodes. Black arrows on images illustrate brace root primordia. B) Distinct transcriptional signatures were identified across the developmental stages. Heatmap scale indicates Z-score relative expression within each gene. The black box delineates genes that were expressed highest in nodes with no primordia and reduced in nodes with primordia, including six SBP transcription factors. **ALT:** (A) Photograph of maize stem and associated imaging of the cross sections of three aboveground nodes of V6 maize plants. Image highlights the three successive stages of brace root development: no primordia, primordia formation, and emergence. (B) Heatmaps showing differential gene expression across brace root developmental stages, with an enlarged section highlighting six SBP transcription factors. SBP transcription factors show distinct expression patterns across brace root stages. Colors illustrate relative expression with red indicating higher and blue indicating lower expression.

Based on the morphological findings, the three aboveground nodes closest to the soil from the V6 stage were subjected to whole-node transcriptome analysis to identify regulators of brace root development. Although whole nodes were analyzed, each node exhibited a distinct transcriptome profile (**Fig S2**; **Fig 1B**), lending credibility to the idea that regulators of brace root development can be identified from this dataset. We hypothesized that differentially expressed genes (DEGs) between nodes with no primordia and nodes with primordia are candidate regulators of initiation, and DEGs between nodes with primordia and emerging primordia are candidate regulators of emergence. For initiation, there were 2,977 DEGs, with 1,896 upregulated (candidate positive regulators) and 1,081 downregulated (candidate negative regulators) (**Table S2 & S3**). For emergence, there were 1,781 DEGs, with 1,156 upregulated and 625 downregulated **(Table S2 & S4**).

To identify DEGs that might regulate different stages of brace root development, transcription factors (TFs) were prioritized. Among the most downregulated TFs during brace root initiation were six SBP TFs, which were expressed highest in nodes with no primordia and reduced in nodes with primordia (**Fig. 1B**). Notably, the SBP TF that was previously implicated in brace root development, TGA1 (Wang et al., 2015), was not differentially expressed in our analysis. However, three of the six differentially expressed SBP TFs (*tsh4*, *ub2*, and *ub3*) were previously implicated in lateral organ branch number. Therefore, we hypothesized that these TFs also play a role in brace root initiation.

### Brace root initiation is increased in *sbp* mutants

Expression patterns suggested that SBP TFs act as negative regulators of brace root initiation. Thus, we predicted that *sbp* mutants would exhibit increased brace root initiation compared to wild-type plants. Increased initiation could result in either more nodes that produce brace roots or more brace roots per node. The number of nodes that produced brace roots (emerged and pre-emerged) was quantified in wild-type (W22) and mutant (*tsh4*, *ub3*, *ub2;ub3*, and *ub2;ub3;tsh4*) plants. W22 plants averaged 4.5 nodes with brace roots, whereas *tsh4* averaged 4.9, *ub3* averaged 5.7, *ub2;ub3* averaged 6.1, and *ub2;ub3;tsh4* averaged 7.3 nodes (**Fig 2A-B**). Thus, the loss of SBP TFs led to a dose-dependent increase in the number of nodes with brace roots.

**Fig 2.**
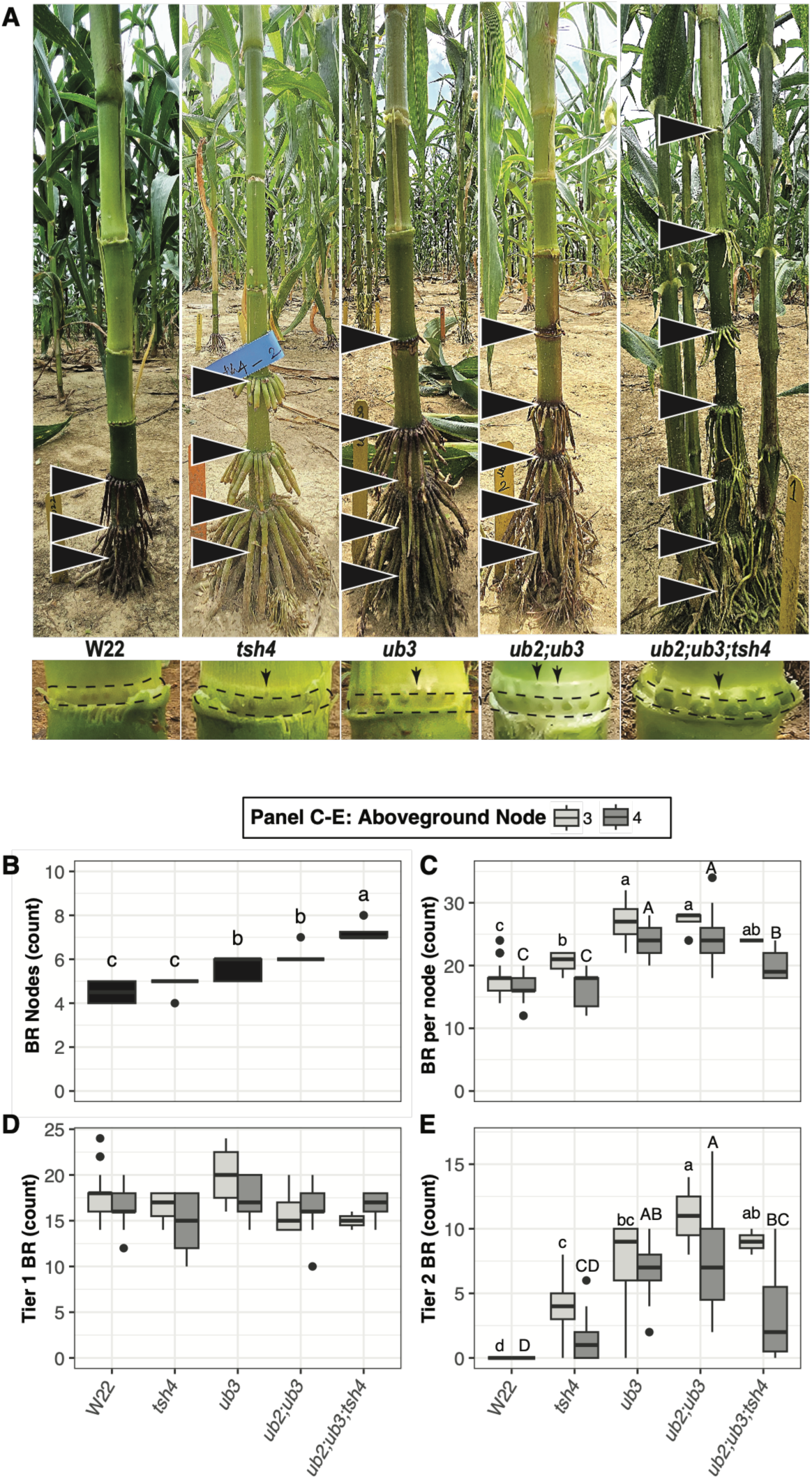
*sbp* mutants increase the number of nodes with brace roots and the number of brace roots within a node by forming a second tier of brace roots. A) *sbp* mutants showed a dose-dependent increase in the number of nodes with brace roots (BR). Arrowheads indicate a node with brace roots. At the node, all genotypes had one tier of brace roots (black dashed outline), and mutants showed variable extent of a second tier of brace roots (arrowheads). B) The total number of nodes containing brace roots increased in *sbp* mutants in a dose-dependent manner. C) The total number of brace roots on aboveground nodes 3 and 4 increased in *sbp* mutants in a non-linear dose-dependent manner. D) The total number of brace roots on tier 1 for aboveground nodes 3 and 4 was similar across all genotypes. E) There were no tier 2 brace roots identified in W22 plants, whereas the *sbp* mutants showed a non-linear dose-dependent number of second tier of brace roots for aboveground nodes 3 and 4. (B-E) Brace roots were considered if primordia or emerged roots were present. Genotypes that share a letter (both case and letter) within a phenotype are not statistically different from one another (P > 0.05). Aboveground nodes 3 and 4 were analyzed separately. Black dots indicate outliers. **ALT:** (A) Photographs of field grown maize plants. The genotypes (W22, *tsh4*, *ub3*, *ub2;ub3*, *ub2;ub3;tsh4*) show differences in the number of nodes producing brace roots. Black arrowheads identify brace root nodes in images. (B–E) Boxplots comparing the number of nodes with brace roots, total brace roots per node, and brace roots in the lower and upper tiers in both aboveground node 3 and 4. Mutants generally have more nodes producing brace roots and more brace roots in the second tier). Box colors illustrate node being assessed.

We additionally assessed the number of brace roots at two nodes – the third aboveground node (node 3) and the fourth aboveground node (node 4), where node 3 is at a later stage of development than node 4. For node 3, W22 averaged 17.8 brace roots per node, whereas *tsh4* averaged 20.5, *ub3* averaged 27, *ub2;ub3* averaged 27, and *ub2;ub3;tsh4* averaged 24 brace roots per node (**Fig 2C**). For node 4, W22 averaged 16.7 brace roots per node, whereas *tsh4* averaged 16.2, *ub3* averaged 24, *ub2;ub3* averaged 24.2, and *ub2;ub3;tsh4* averaged 20 brace roots per node (**Fig 2C**). There was a non-linear dose-dependent increase in the number of brace roots in both nodes. This was observed as an increase in brace roots in the single and double mutants that was attenuated in the triple mutant. Together, these results show that SBP TFs limit both the number of nodes that produce brace roots and the number of brace roots per node.

We were interested in understanding the basis of brace root number increases. In wild-type maize, brace roots form in a single concentric tier around the circumference of the node and are therefore geometrically restricted. To accommodate more brace roots, the node diameter must increase, or the roots must become thinner. However, *sbp* mutants displayed a third option and developed two tiers of brace roots at each node (**Fig 2A**). We quantified the number of brace roots in each tier for node 3 and node 4. For the lower tier 1, there were no statistical differences within a node between W22 plants (average 17.8 and 16.7 brace roots in node 3 and 4, respectively), *tsh4* (16.5 and 17.4), *ub3* (20 and 17.4), *ub2;ub3* (16 and 16.6), and *ub2;ub3;tsh4* (15 and 16.6) (**Fig 2D**). W22 never developed a second tier of brace roots, whereas *tsh4* averaged 4 and 1.5 (node 3 and node 4, respectively), *ub3* averaged 7 and 6.6, *ub2;ub3* averaged 11 and 7.9, and *ub2;ub3;tsh4* averaged 9 and 3.4 (**Fig 2E**). No second tier of brace roots were observed in *Cg1* mutants (**Fig S3**), demonstrating that the extent of second tier occupancy displays a non-linear dose-dependent response consistent with the increased brace root number.

Developmental programs that regulate brace roots have been posited to be generalizable to crown root development. However, when the *sbp* mutants were assessed for crown root tier expansion, there were no tier 2 crown roots identified (**Fig S4**). This observation suggests that despite both root types developing from the stem node, there may be divergent pathways controlling brace root and crown root development.

### SBP TFs limit the size and competency of the stem pulvinus

A second tier of brace root primordia could result from node duplication, expansion of the competence zone (i.e., pulvinus), and/or de-repression of brace root initiation programs. Node duplication was ruled out because other nodal organs, such as the leaf or axillary meristem, were not duplicated. To determine whether an expansion of the competence zone was associated with a second-tier formation, the length of the pulvinus was measured (**Fig 3A**). Consistent with the non-linear dosage in tier 2 brace root primordia, pulvinus length also increased non-linearly (**Fig 3B**). However, pulvinus length was poorly correlated with the number of brace roots in tier 2 (**Fig 3C**). This indicates that the expansion of the competence zone is not sufficient for the formation of a second tier of brace root primordia and supports a model in which SPB TFs actively repress brace root initiation.

**Fig 3.**
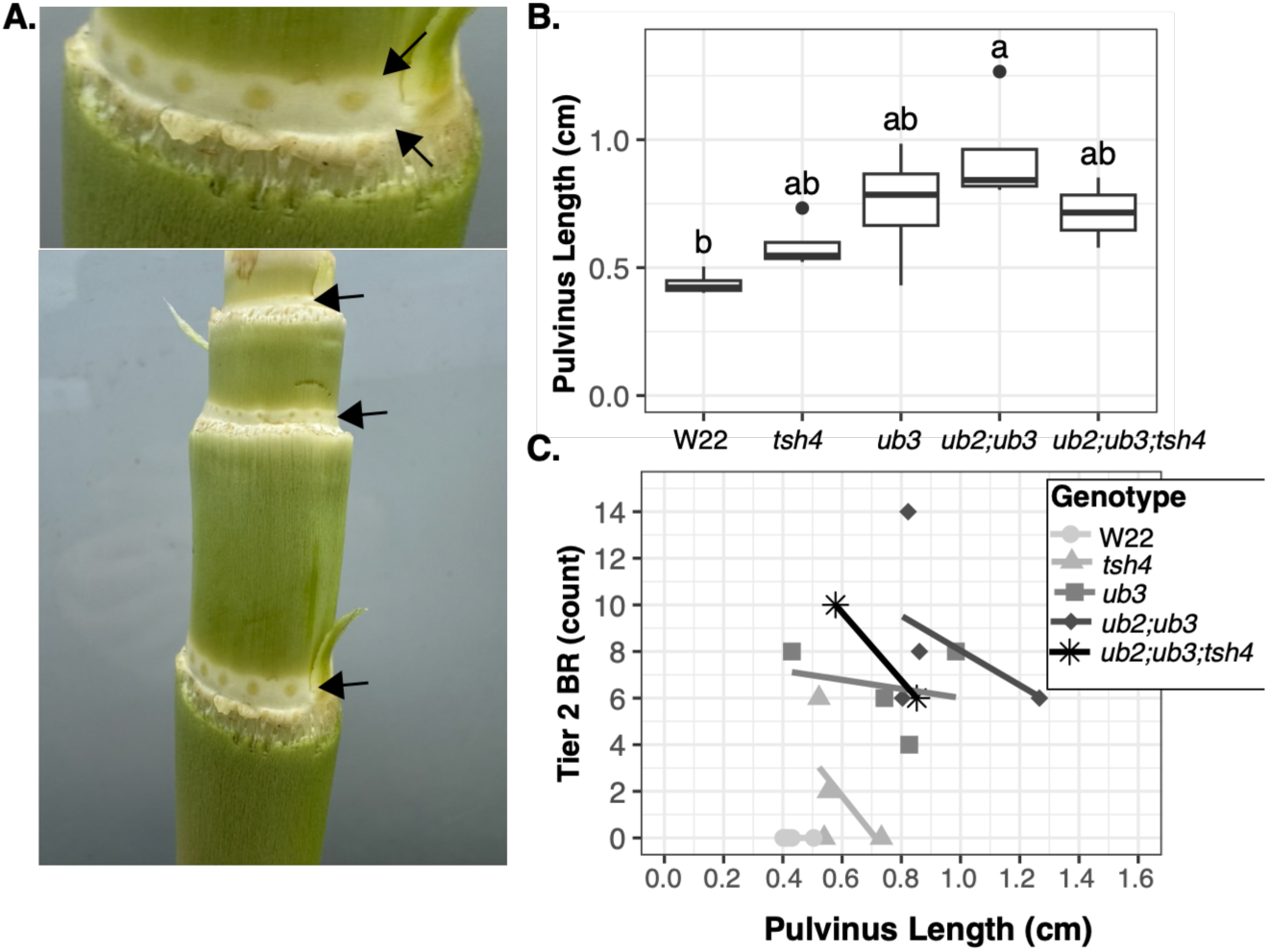
*sbp* mutants have an increased pulvinus length. A) The stem pulvinus is directly above the leaf sheath attachment. It is visible as the lighter colored band as highlighted with black arrows. B) The pulvinus length increased non-linearly in *sbp* mutants, with *ub2;ub3* being the only genotype statistically different from W22 (P < 0.05). Genotypes that share a letter are not statistically different from one another. Black dots indicate outliers. C) The pulvinus length was poorly correlated with the number of brace roots (BR) in the second tier. **ALT:** A) Photographs showing the maize stem pulvinus and brace root primordia. Arrows highlight the stem pulvinus. (B) Boxplots comparing pulvinus length across W22 and *sbp* mutant genotypes. (C) Scatterplot showing the relationship between pulvinus length and tier 2 brace root number, with genotype-specific regression lines.

### SBP regulatory networks during brace root initiation are distinct from DNA binding targets in immature ears

TSH4, UB2, and UB3 have been previously reported to promote inflorescence branch number (Chuck et al. 2010). Therefore, we explored the hypothesis that SBP regulatory networks are rewired to inhibit brace root initiation. In a rewired network, downstream SBP targets in inflorescence would overlap with brace root initiation candidates. To test this, downstream DNA binding targets of TSH4 and UB3 from immature ears were intersected by combining TSH4 Chromatin IP (ChIP)-seq (Dong et al. 2024) and UB3 DNA Affinity Purification (DAP)-seq (Ricci et al. 2019; Dong et al. 2024) data. This overlap identified 886 shared downstream DNA binding targets. These downstream targets were then intersected with brace root initiation candidates, identifying 157 overlapping genes (87 downregulated and 70 upregulated during brace root initiation) (**Table S5**). We rationalized that if these genes function downstream of SBP TFs during brace root initiation, their expression would also be altered in *sbp* mutant nodes. Since *sbp* mutants exhibit greater brace root initiation (**Fig 2**), genes upregulated during initiation would have increased expression in mutants, whereas genes downregulated during initiation would have reduced expression in mutants. To assess this regulation, the second aboveground node from *tsh4* and *ub2;ub3* mutant plants was dissected into two tiers (T1 and T2) for transcriptome profiling. Because UB2 and UB3 are duplicated in maize (Chuck et al. 2014), the *ub2;ub3* mutant was used. Each mutant tier (*tsh4-*T1, *tsh4*-T2, *ub2;ub3*-T1, *ub2;ub3*-T2) transcriptome was compared to the wild type (W22) tier 1 transcriptome **(Table S2, S6-S9**). There were 1,678 DEGs that were common in all four mutant-vs-wild type comparisons (**Table S10**). Among these DEGs, only 12 genes were identified as SBP targets in immature ears, and only four of these genes changed in a direction consistent with SBP regulation (**Table S11**). These findings suggest that the regulatory networks controlled by SBP TFs during brace root initiation are largely distinct from those occurring in the ear.

### SBP TFs limit brace root initiation by inhibiting *liguleless2*

To investigate the regulatory networks controlled by SBP TFs during brace root initiation, we revisited the list of the DEGs shared in all four comparisons (**Table S10**) and checked for overlap with the brace root initiation candidates. A total of 223 genes were shared between mutant DEGs and brace root initiation candidates. This list was again refined by direction of gene expression change that would be consistent with SBP regulation. This analysis identified 67 DEGs (35 downregulated and 32 upregulated) that are high-confidence downstream targets of SBP TFs during brace root initiation (**Table S12**). Most of these genes have not been characterized, but a few have previously reported roles in regulating cell division and cellular processes. For example, genes involved in DNA replication or cell division (e.g., *minichromosome maintenance 2*; *indeterminate gametophyte 3*) were downregulated; a gene from a family known to regulate plant and organ size (*cell number regulator 7* (Guo et al. 2010)) was upregulated during initiation and in *sbp* mutants. Only one gene with a known role in phytohormone signaling was identified; *abscisic acid stress ripening1* (*aasr1*) was upregulated during brace root initiation and in *sbp* mutants. *aasr1* is upregulated under drought and abscisic acid (ABA) treatment and negatively regulates drought tolerance (Yang et al. 2024).

Among these candidates, *liguleless2* (*lg2*) was investigated further based on its previously characterized role in maize development and its expression patterns, which were consistent with SBP-mediated regulation. Specifically, *lg2* was upregulated during brace root initiation and in *sbp* mutants (**Fig 4A-B**). *lg2* has been previously reported to coordinate with *liguleless1* (*lg1*) to establish the leaf ligule and auricle boundaries (Harper and Freeling 1996). Both *lg1* and *lg2* were upregulated during brace root initiation (**Fig 4A**), suggesting that these genes act as a positive regulator of brace root initiation. However, these genes exhibited distinct expression patterns in *sbp* mutants, with *lg2* significantly upregulated in all four mutant comparisons, whereas *lg1* was significantly downregulated only in *tsh4* tier 1 (**Fig 4B**). When assessing phenotypic differences in *lg1* and *lg2* mutant plants, we found that the single *lg1* and *lg2* mutants did not alter the number of nodes with brace roots in the field (**Fig 4C**), and the *lg2* mutant in the greenhouse did not alter the number of roots within a node (**Fig 4D**). However, the double *lg1;lg2* mutant produced no nodes with brace roots (**Fig 4C**). These results validate the high confidence gene list, while also emphasizing the redundancy of *lg1* and *lg2* for brace root initiation. Together, these data demonstrate that SBP TFs inhibit brace root initiation through the activation of *lg2*, which functions in concert with *lg1*.

**Fig 4.**
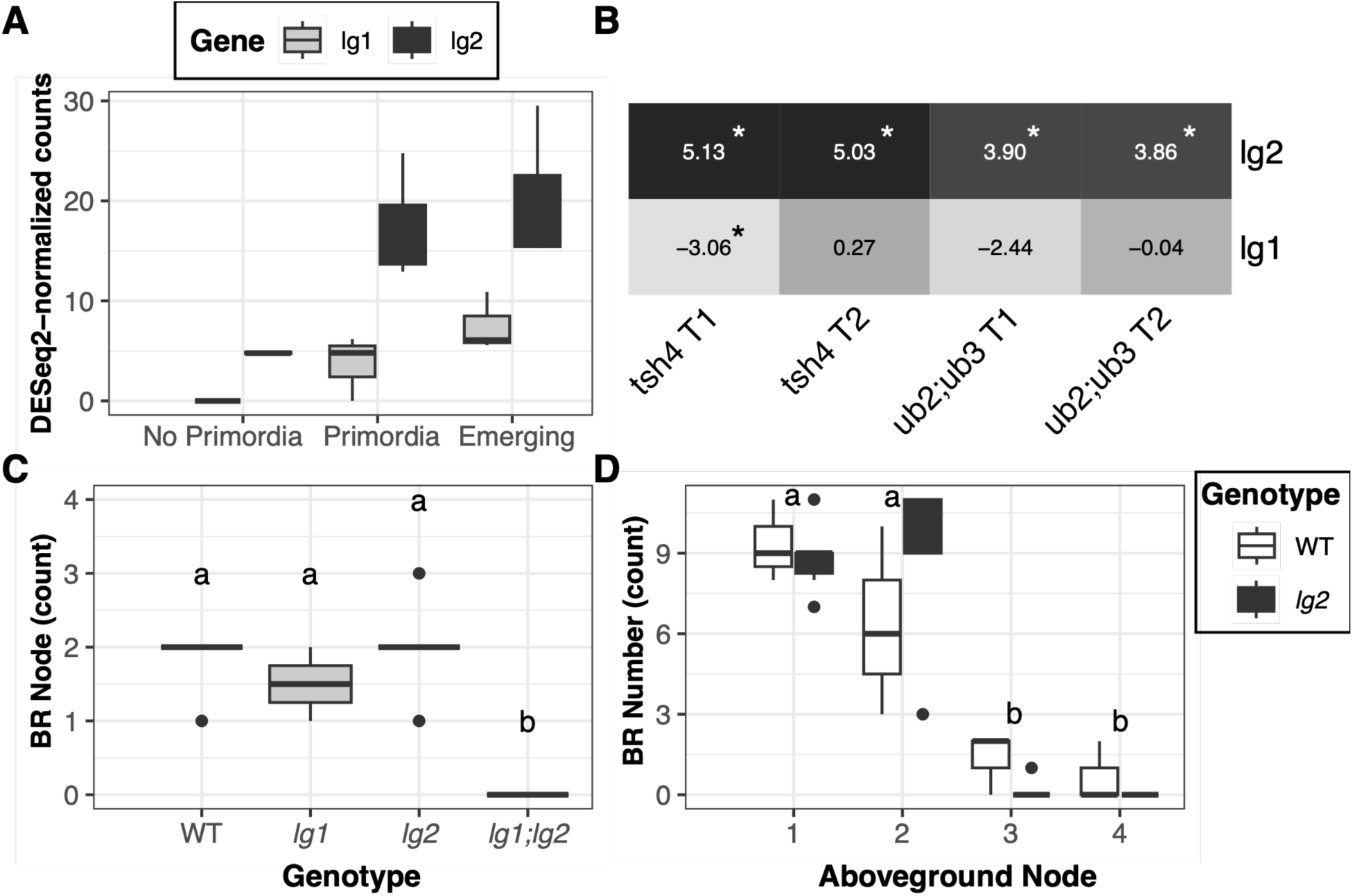
*lg1* and *lg2* function together to inhibit brace root initiation. A) The expression of *lg1* and *lg2* increased across nodes with brace root (BR) primordia and emerging primordia. B) *lg2* was significantly increased in in *tsh4* and *ub2;ub3* mutants (tier 1 and 2) compared to wild-type (tier 1) plants. In contrast, *lg1* expression was decreased in mutant *tsh4* tier 1 and *ub2;ub3* tier 1 compared to wild-type (tier 1) plants, although only the *tsh4* comparison was statistically significant. Positive values indicate increased expression in mutants relative to the wild type, whereas negative values indicate decreased expression in mutants relative to the wild type. Asterisks indicate statistically significant differential expression. T# indicates tier (T1 or T2). C) There were fewer nodes containing brace roots in the double mutant (*lg1;lg2*) compared to the wild type or single mutant (*lg1* or *lg2*). Genotypes that share a letter are not statistically different from one another (P > 0.05). D) The total number of brace roots on each node (aboveground nodes 1-4) were similar between *lg2* and its wild-type sibling (P > 0.05). Nodes that share a letter are not significantly different from one another (P > 0.05). (C-D) Black dots indicate outliers. **ALT:** (A) Boxplots showing *lg1* and *lg2* expression across three brace root developmental stages (no primordia, primordia formation, and emergence). The color of the boxplot illustrates gene (lg1 or lg2) (B) Heatmap showing differential expression of both lg1 and lg2 in both the lower and upper tiers of *tsh4* and *ub2;ub3* mutants, with numerical log2 fold changes. (C) Boxplots comparing the number of nodes containing brace roots in the wild type and mutants (*lg1*, *lg2*, and *lg1;lg2*). (D) Boxplots showing the number of brace roots across the four aboveground nodes closest to the soil in wild type and *lg2* plants.

### SBP TFs inhibit brace root initiation through downstream regulation of ACC

Although only one phytohormone-related gene was in our high confidence SBP target list, we anticipate that phytohormones play an important role in the regulation of brace root initiation. To further define the pathways downstream of SBP TFs, phytohormones – ABA, the ethylene precursor 1-aminocyclopropane-1-carboxylic acid (ACC), intermediate forms of gibberellin precursors (GA_12_ and GA_53_), the bioactive gibberellin (GA_4_), indole-3-acetic acid (IAA; the most prevalent form of auxin), 12-hydroxyjasmonic acid (12-OHJA; an inactive JA derivative), jasmonic acid (JA), and salicylic acid (SA) – were quantified in tier 1 tissue from the second aboveground node of W22, *tsh4*, and *ub2;ub3* plants. All hormones were detected except IAA, which was below the detection limit in all samples. This analysis showed that ABA, GA_53_, and JA did not differ significantly among genotypes (**Fig 5**). ACC, the ethylene precursor, was the only hormone significantly elevated in both *tsh4* and *ub2;ub3* relative to W22 (**Fig 5**). This finding is consistent with previous reports showing that ACC levels are positively correlated with brace root node number (Wei et al. 2026), and that ethylene promotes brace root emergence (Shi et al. 2019). We anticipated more similarities than differences between the two mutants, but we instead identified many differences in phytohormone profiles between *tsh4* and *ub2;ub3*. Specifically, GA_12_ was lower in *tsh4* and the active GA_4_ was lower in *ub2;ub3* (**Fig 5**). Whereas SA and 12-OHJA were higher in *ub2;ub3* (**Fig 5**). Together, these data highlight a shared role of SBP TFs in regulating ACC accumulation and identify differences in downstream regulation of phytohormones by TSH4 and UB2/UB3.

**Fig 5.**
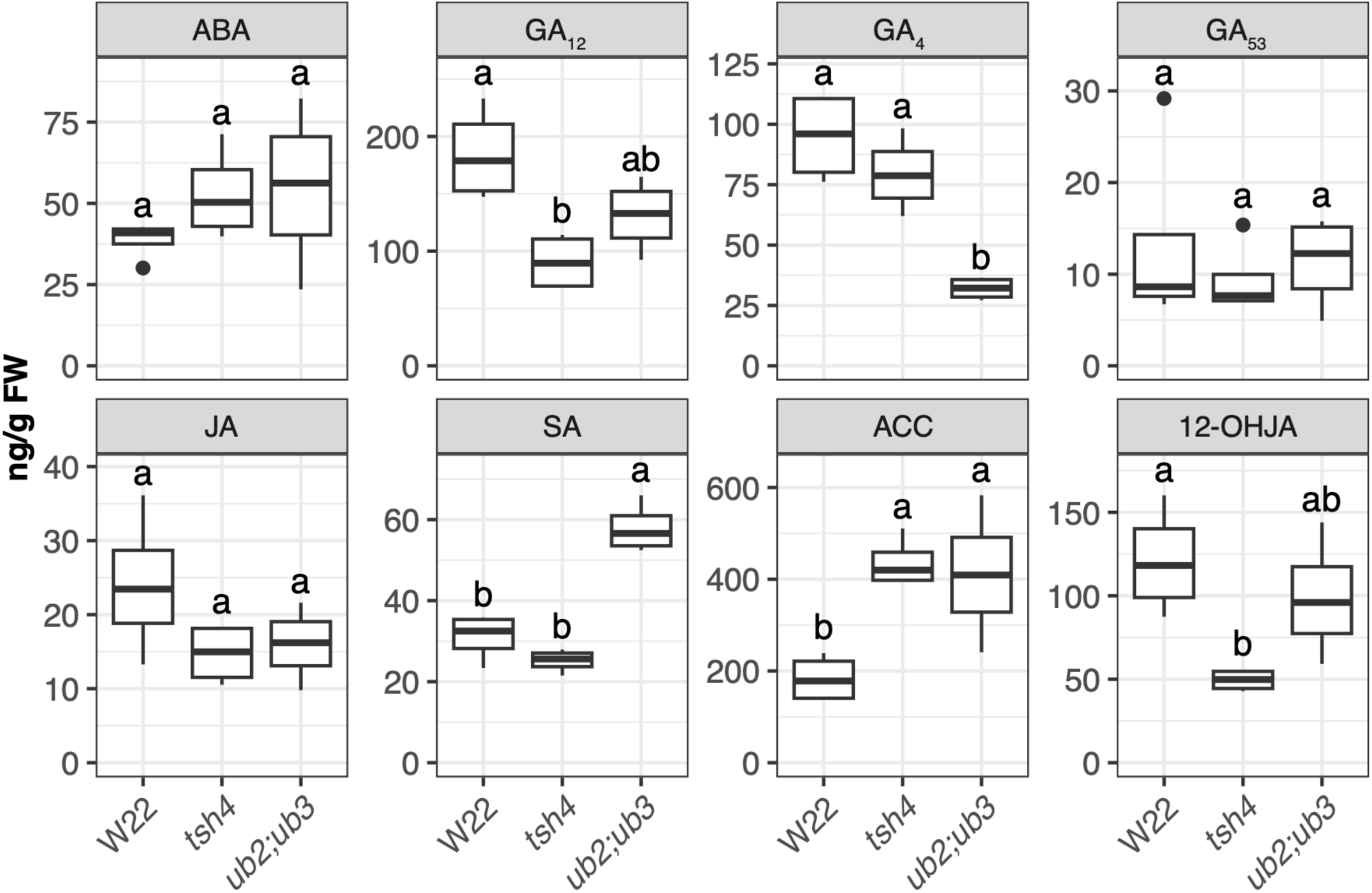
*tsh4* and *ub2;ub3* have both shared and divergent phytohormone profiles. ACC, the precursor of ethylene was elevated in nodes of *tsh4* and *ub2;ub3* mutants compared to W22. In contrast, SA was elevated in *ub2;ub3* mutants, and GA_4_ was reduced in *ub2;ub3* mutants. All other phytohormones showed no significant changes among genotypes. Genotypes that share a letter are not statistically different from one another (P > 0.05). Black dots indicate outliers. **ALT:** Eight sets of boxplots comparing ABA, GA_12_, GA_4_, GA_53_, JA, SA, ACC, and 12-OHJA in W22, *tsh4*, and *ub2;ub3* plants. GA_4_ is reduced in *ub2;ub3*, while ACC is elevated in both mutants relative to W22. Letters indicate statistical groupings.

### Gibberellin promotes brace root growth after initiation

To explore the differences in downstream regulation of brace root initiation between TSH4 and UB2;UB3, T1 transcriptomes were compared between *tsh4* and *ub2;ub3* mutants and these DEGs were intersected with brace root initiation candidates (**Table S13**). Among the 393 DEG between the mutants that were initiation candidates, 220 had higher expression in *tsh4* and 173 had higher expression in *ub2;ub3* (**Table S13**). Among these DEGs, the gibberellin biosynthesis gene *d3* was upregulated 2.85-fold in *tsh4* relative to *ub2;ub3*, consistent with the levels of the bioactive gibberellin (GA_4_) being higher in *tsh4* compared to *ub2;ub3* (**Fig 5**).

Gibberellin typically promotes organ growth by inducing cell division and elongation (Robil et al. 2024). Therefore, we hypothesized that the differential expression of gibberellin would lead to different brace root phenotypes. Indeed, we found that *tsh4* plants produced fewer, but larger brace roots compared to *ub2;ub3* plants, which had more, thinner brace roots (**Fig 6A-B**). These findings support a model in which *tsh4* and *ub2;ub3* act antagonistically through gibberellin to promote brace root growth after initiation (**Fig 6C**). To further test the role of GA in brace root initiation versus post-initiation growth, brace root phenotypes were quantified in *d3* mutant plants and their wild-type siblings. There was no difference in stem diameter, brace root size, or brace root number between mutants and wild-type siblings (**Fig S5**). However, there was also no increase in the number of nodes that produced brace roots, contrary to previous report for the *d3* mutant in the A632 background (Evans and Poethig 1995). The varying phenotypes are likely due to the background-specific interactions, which have been previously shown to impact gibberellin responses (Best and Dilkes 2022). The background-specific interaction in the context of brace root initiation is supported by the triple *ub2;ub3;tsh4* having more nodes that produced brace roots in the B73 background compared to the W22 background (**Fig S6**). Thus, the mechanism by which *tsh4* and *ub2;ub3* act antagonistically through gibberellin to promote brace root growth after initiation may be background dependent.

**Fig 6.**
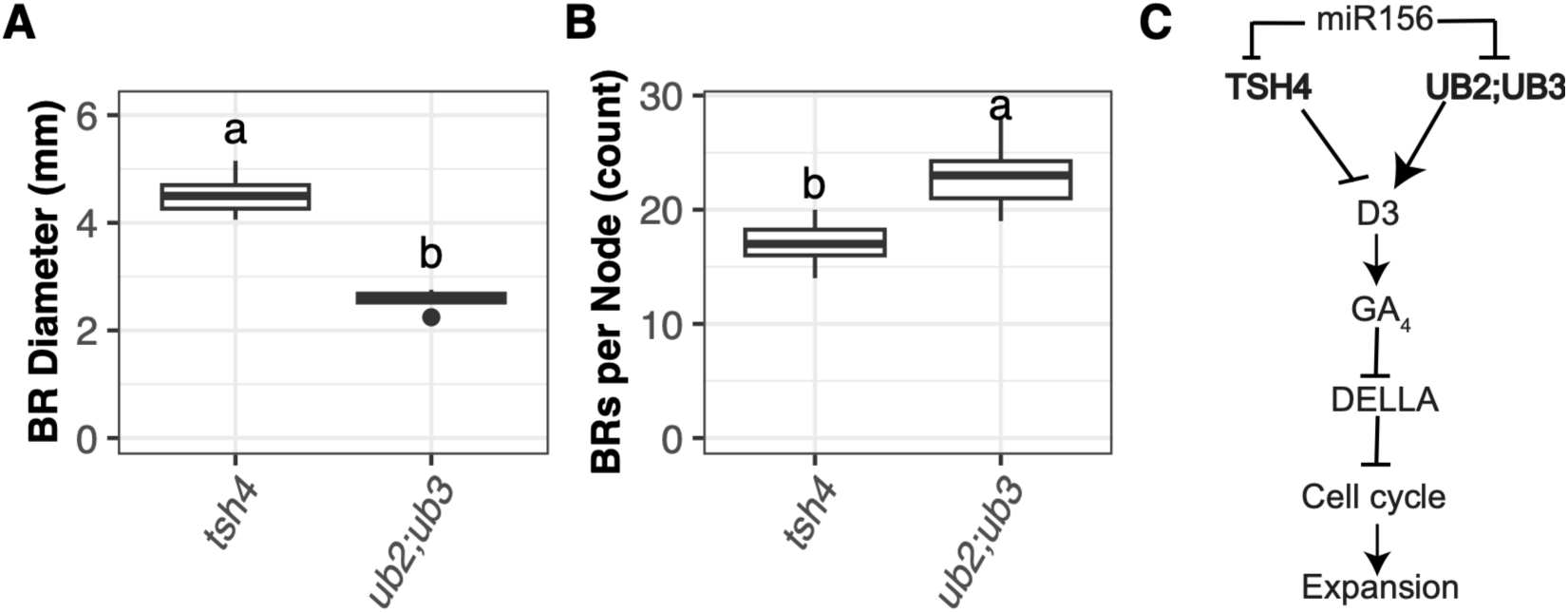
TSH4 and UB2;UB3 have differential brace root phenotypes established through gibberellin-mediated primordia expansion. A) Emerged brace roots (BRs) of *tsh4* were larger in diameter than those of *ub2;ub3*. B) *ub2;ub3* had more brace roots per node than *tsh4*. C) The model for antagonistic establishment of brace root size with TSH4 inhibiting *d3* expression, and UB2;UB3 promoting *d3* expression to induce primordia expansion. **ALT:** (A–B) Boxplots comparing brace root diameter and number of brace roots per node in *tsh4* and *ub2;ub3* mutants. *tsh4* produces fewer, thicker roots, whereas *ub2;ub3* produces more, thinner roots. (C) Proposed regulatory model connecting miR156, TSH4 and UB2;UB3, D3, gibberellin signaling, DELLA, and cell cycle regulation to brace root development. Arrows indicate positive regulation and blunt-ended lines indicate repression.

### A balance between *ub3* and *tsh4* expression underlies natural variation in brace root size

The contrasting phenotypes of *tsh4* and *ub2;ub3* mutant plants (**Fig 7A**) suggest that the relative activity of SBP TFs within a node influences the balance between brace root initiation and subsequent growth. To determine if the relative activity of *tsh4* and *ub3* expression contributes to variation in brace root phenotypes in wild-type germplasm, or if this is also background-dependent, we leveraged data from a previous study that quantified brace root phenotypes in 52 maize inbred lines (Hostetler et al. 2022). Inbred lines were visualized as a scatterplot of brace root diameter versus brace root number (**Fig S7**). From this visualization, CML333 was selected as a genotype that had more, thinner brace roots, and PHHB4 was selected as a genotype that had fewer, thicker brace roots, with both lines having statistically similar stem widths (**Fig 7B**; **Fig S7**).

**Fig 7.**
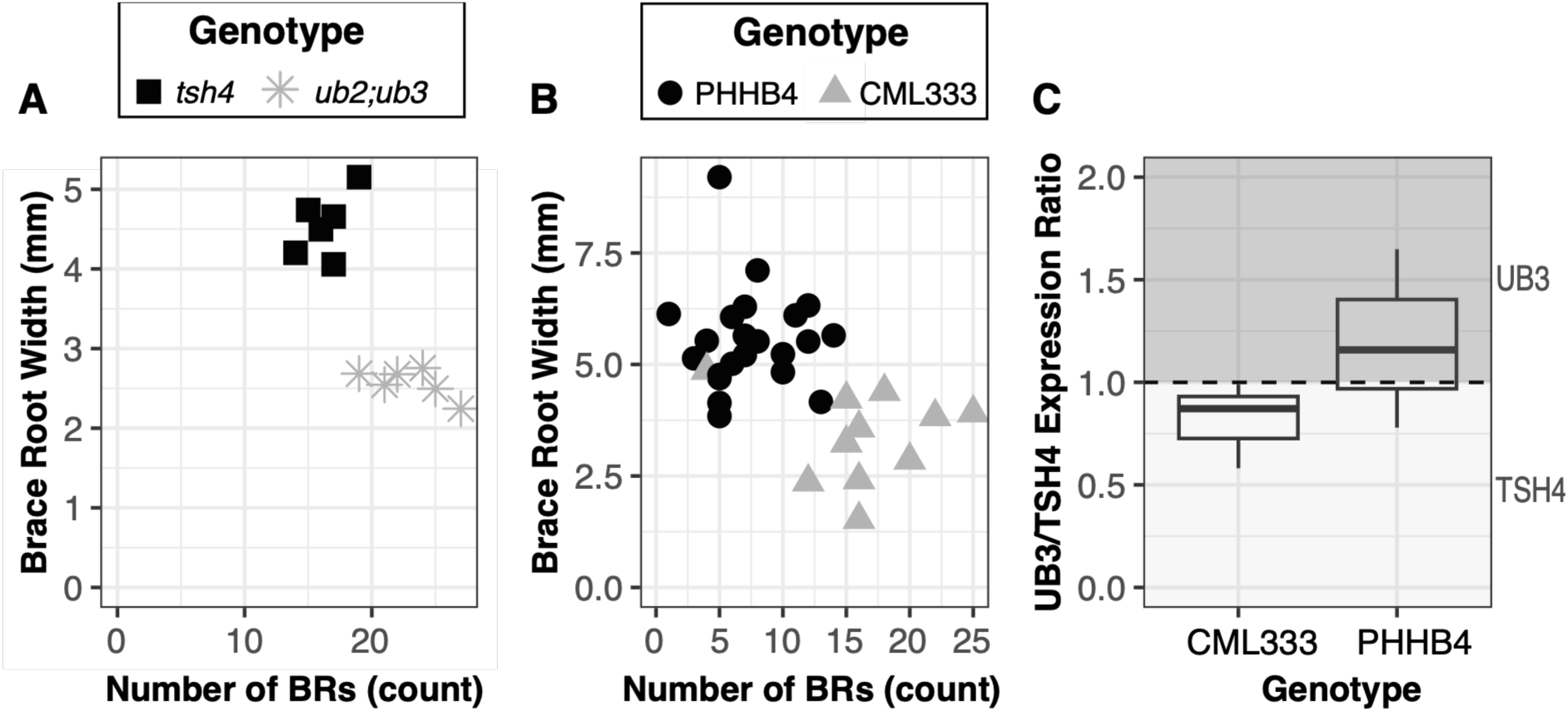
*tsh4* and *ub2;ub3* antagonism underlies natural variation in brace root size and number. A) *tsh4* had fewer thicker brace roots compared to *ub2;ub3*, which had more thinner brace roots. Genotypes did not statistically differ for stem width (P>0.05). B) The inbred line, PHHB4, had fewer thicker brace roots (like *tsh4*) compared to CML333, which had more thinner brace roots (like *ub2;ub3*). Genotypes did not statistically differ for stem width (P>0.05). C) The relative UB3/TSH4 transcript ratio in CML333 and PHHB4 shows that the UB3/TSH4 is approximately 47% higher in PHHB4 compared to CML333. The dotted line at 1 indicates equal expression of *ub3* and *tsh4*. Values above 1 (dark gray region) indicate higher *ub3* expression relative to *tsh4,* whereas values below the dotted line (light gray region) indicate lower *ub3* expression relative to *tsh4*. **ALT:** (A–B) Scatterplots comparing brace root number and size in (A) *tsh4* and *ub2;ub3,* and (B) maize inbred lines CML333 and PHHB4. Genotypes differ in the relationship between root number and size, with *tsh4* and PHHB4 having fewer thicker brace roots, while *ub2;ub3* have more thinner brace roots. (C) Boxplots showing the UB3-to-TSH4 expression ratio in CML333 and PHHB4, with a higher ratio in PHHB4.

To assess whether the balance of *sbp* expression corresponded to these contrasting phenotypes, TSH4 and UB3 transcript abundance was quantified in both CML333 and PHHB4. We hypothesized that if the relative balance of *sbp* gene expression influences brace root architecture, then CML333, which produces more thinner roots, would have decreased UB3 relative to TSH4 and PHHB4, which produces fewer thicker roots, would have increased UB3 relative to TSH4. Consistent with this expectation, the expression ratio of UB3/TSH4 is about 47% higher in PHHB4 compared to CML333 (**Fig 7C**). These results demonstrate that the balance of TSH4 and UB3 expression contributes to natural variation in brace root size and number.

## Discussion

Although nodal roots are critical for maize production, our understanding of their developmental regulation is limited. In this study, transcriptome profiling across brace root developmental stages revealed a finely tuned, stage-specific regulatory framework, with *tsh4*, *ub2,* and *ub3* identified as key repressors of brace root initiation. The *tsh4*, *ub2* and *ub3* mutants have been previously characterized for their role in inflorescence development with *tsh4* and *ub2/ub3* independently promoting tassel branch number, *ub2/ub3* limiting kernel row number, and *tsh4* repressing bracts (Chuck et al. 2010, 2014); while all three redundantly repress tillers (Chuck et al. 2014; Liu et al. 2021). These results highlight the differential organogenesis programs downstream of SBP TFs. We initially hypothesized that inflorescence organogenesis programs would share downstream targets that are re-wired during brace root initiation. However, the minimal overlap in gene targets suggests that these programs are regulated through distinct downstream mechanisms that are shaped by developmental context. Indeed, in ears, SBP TFs predominantly engage auxin-mediated pathways through direct regulation of repressor AUX/IAA genes (Kong et al. 2023; Dong et al. 2024). In this study, auxin accumulation was not detected in any of the samples analyzed, and no AUX/IAA genes were identified among the initiation candidates and differentially expressed genes in *sbp* mutant nodes. In contrast, we show that brace root initiation relies on SBP-mediated de-repression of ethylene to promote cell cycle progression and organ initiation. This is consistent with previous reports that have emphasized the importance of ethylene in brace root development (Shi et al. 2019; Wei et al. 2026).

We anticipated that SBP TFs function redundantly to repress brace root initiation based on the dose-dependent increase in brace root number following reductions in SBP activity. Instead, we identified distinct downstream pathways of *tsh4* and *ub2;ub3*, which modulated brace root number and size. Specifically, gibberellin levels were reduced in *ub2;ub3* mutants but remained unchanged in *tsh4* mutants relative to the wild type, suggesting that UB2;UB3 regulates brace root development through gibberellin-associated pathways. This differs from previous reports in maize bract primordia which showed that TSH4 suppresses gibberellin to suppress bract growth (Xiao et al. 2022). Furthermore, an analysis of two inbred maize lines with contrasting brace root occupancy strategies (fewer, thicker versus more, thinner), showed that differences in *tsh4* and *ub2*;*ub3* expression is consistent with these pathways regulating natural variation in brace root phenotypes. These data further support a new model of nodal root development that includes a stage that regulates primordia expansion before emergence.

A predictive network of bioenergy *Sorghum bicolor* (sorghum) brace root development had previously identified the *Zm*TSH4 ortholog *Sb*SPL9 as a central regulator within a brace root gene regulatory network (Lamb et al. 2024). However, this network had *Sb*SPL9 functioning as a positive regulator of initiation, whereas we show that *Zm*TSH4 repress primordia initiation. Bioenergy sorghum also produces two brace root tiers per node, but in contrast to the maize *sbp* mutants, the brace roots do not emerge from the second tier (Lamb et al. 2024). Thus, while bioenergy sorghum and maize may share regulatory components that control brace root initiation, these components may mediate different developmental outcomes.

The results presented here show that *sbp* mutants impact brace root initiation but not crown root initiation. One of the possible explanations for this may lie in the stem pulvinus. The pulvinus, defined as an aboveground stem feature (Brown et al. 1959), is the nodal structure from which brace root primordia arise, and our data show that SBP dosage directly controls both its physical size and the initiation programs within it. In contrast, crown root nodes are subterranean, highly compressed, and have no reported pulvinus. This means that the nodal structure within which SBP-regulated programs act is likely absent. Together, these findings indicate that brace root initiation requires the de-repression of aboveground-specific SBP transcriptional programs and their downstream targets.

This study fills a gap in the molecular regulation of brace root development, demonstrating the ability to modulate development through integrated TF and hormonal signaling. However, SBP TFs function broadly across multiple organs, and due to this pleiotropy, tissue specific repression of these TFs would be necessary for selectively modulating brace root traits. Validating tissue-specific regulators downstream of SBP TFs is necessary for precise crop improvement strategies. In this study, we verified the role of *lg1* and *lg2* to promote brace root initiation. This is particularly interesting given that there has been selection on *lg1* and *lg2* variants that promote more upright leaf architecture and support higher planting densities (Ding et al. 2015; Sarma et al. 2026). This suggests that selection for leaf angle may indirectly select for reduced brace roots. Continued dissection of these regulatory pathways and an understanding of the pleiotropic effects of crop improvement targets will be critical to fully elucidate and precisely engineer maize root systems.

## Acknowledgements

We thank Sparks Lab members, Dr. Paula McSteen (University of Missouri), and Dr. Elizabeth Haswell (The Sustainable Professor) who provided feedback on this study, and Nate Harlan for growing plants and collecting initial data; Dr. Jingjing Tong for help with the paper germination experiment. We additionally acknowledge Dr. George Chuck for providing *sbp* mutant seed; Dr. Paula McSteen and Dr. Prameela Awale for providing *d3* mutant seed; Dr. Madelaine Bartlett, Dr. Amanda Rasmussen and Dr. Jesus Beltran for helpful discussion during thesis committee meetings for TK. RNA-sequencing was completed at the University of Delaware DNA Sequencing & Genotyping Center. Mention of trade names or commercial products in this publication was solely for the purpose of providing specific information and does not imply recommendation or endorsement by the U.S. Department of Agriculture.

## CRediT

**Thanduanlung Kamei** - Formal Analysis, Investigation, Writing - original draft, Writing - review & editing; **Ashley N. Hostetler** – Validation, Formal Analysis, Data Curation, Visualization, Writing - original draft, Writing - review & editing; **Sarah Tallents** - Formal Analysis, Investigation, Writing - review & editing; **Taran A. Kermani** - Formal Analysis, Investigation, Writing - review & editing; **Ed Bridge** - Investigation, Writing - review & editing; **Annis E. Richardson** - Investigation, Supervision, Funding Acquisition, Writing - review & editing; **Norman B. Best** - Formal Analysis, Investigation, Funding Acquisition, Writing - review & editing; **Erin E. Sparks** - Conceptualization, Methodology, Validation, Formal Analysis, Resources, Funding Acquisition, Investigation, Project Administration, Supervision, Writing - original draft, Writing - review & editing

## Supplementary Material

**Table S1. *tsh4* and *ub3* primers used for qPCR.** Gene IDs are based on the B73 v5 reference genome.

**Table S2. Normalized gene counts.** Normalized gene counts for each sample are listed. For B73 V6 samples the sample name is as follows: B73 N#P#. N# indicates the aboveground node with N1 being the first aboveground node, closest to the soil and N3 being the top aboveground node, furthest from the soil. P# indicates the plant number (biological replicate). For *sbp* mutant plants (*tsh4* and *ub2;ub3*) and W22, the sample name is as follows: Genotype P#S#. Genotype corresponds to either *tsh4, ub2;ub3*; or W22. P# indicates the plant number (biological replicate). T# indicates tier (T1 or T2). Gene IDs are based on the B73 v5 reference genome annotation.

**Table S3. Differentially expressed genes between primordia-containing and non-primordia tissues (Initiation Candidates).** Genes were considered differentially expressed when the adjusted P value was < 0.01 and the absolute log2FC was > 1. Gene IDs are based on the B73 v5 reference genome annotation.

**Table S4. Differentially expressed genes between tissue containing emerged brace roots and tissue containing brace root primordia (Emergence Candidates).** Genes were considered differentially expressed when the adjusted P value was < 0.01 and the absolute log2FC was > 1. Gene IDs are based on the B73 v5 reference genome annotation.

**Table S5. Overlap of initiation candidates from brace root RNA-seq with SBP targets from immature ears.** There were 157 genes that were shared between TSH4 Chromatin IP (ChIP)-seq data from immature ears, UB3 DNA Affinity Purification (DAP)-seq data from immature ears, and differential gene expression analysis between brace root primordia-containing and non-primordia containing tissues. Genes were considered differentially expressed when the adjusted P value was < 0.01 and the absolute log2FC was > 1. Gene IDs are based on the B73 v5 reference genome annotation.

**Table S6. Differentially expressed genes between *tsh4* tier 1 and W22 tier 1.** Genes were considered differentially expressed when the adjusted P value was < 0.01 and the absolute log2FC was > 1. Both B73 v5 and W22 gene IDs are included.

**Table S7. Differentially expressed genes between *tsh4* tier 2 and W22 tier 1.** Genes were considered differentially expressed when the adjusted P value was < 0.01 and the absolute log2FC was > 1. Both B73 v5 and W22 gene IDs are included.

**Table S8. Differentially expressed genes between *ub2;ub3* tier 1 and W22 tier 1.** Genes were considered differentially expressed when the adjusted P value was < 0.01 and the absolute log2FC was > 1. Both B73 v5 and W22 gene IDs are included.

**Table S9. Differentially expressed genes between *ub2;ub3* tier 2 and W22 tier 1.** Genes were considered differentially expressed when the adjusted P value was < 0.01 and the absolute log2FC was > 1. Both B73 v5 and W22 gene IDs are included.

**Table S10. Differentially expressed genes that are shared between all four *sbp* mutant datasets.** The intersection of the four *sbp* mutant differential gene expression datasets. Genes were considered differentially expressed when the adjusted P value was < 0.01 and the absolute log2FC was > 1. Gene IDs are based on the B73 v5 reference genome annotation. Regulation indicates the direction of change in *sbp* mutants.

**Table S11. Genes that are shared between brace root development and immature ears.** The intersection of initiation candidates, four *sbp* mutant differential gene expression datasets, and downstream targets identified from TSH4 ChIP-seq and UB3 DAP-seq datasets from immature ears. Gene IDs are based on the B73 v5 reference genome annotation. Regulation indicates the direction of change in initiation and *sbp* mutants. Genes retained in this list were also present in TSH4 ChIP-seq and UB3 DAP-seq datasets from immature ear datasets.

**Table S12. High confidence genes for brace root development.** The intersection of initiation candidates with four *sbp* mutant DEGs provides a list of high-confidence downstream targets of SBP TFs during brace root development. Gene IDs are based on the B73 v5 reference genome annotation. Regulation indicates the direction of change in initiation and *sbp* mutants.

**Table S13. Differentially expressed genes between tsh4 tier 1 and ub2;ub3 tier 1.** Genes were considered differentially expressed when the adjusted P value was < 0.01 and the absolute log2FC was > 1. Gene IDs are based on the B73 v5 reference genome annotation.

## Funding acknowledgements

This work is supported by the National Science Foundation (NSF), IOS Award #2109189 (EES) and the United States Department of Agriculture-Agricultural Research Service (USDA-ARS), Project #5070-21220-046-000-D (NBB). Funding was provided by the University of Edinburgh Start-Up Fund (A.E.R.), UKRI EP/Y010116/1 (A.E.R.) and Royal Society IES\R2\212032 (A.E.R.)

## Data Availability

All raw data and R scripts used to process and analyze data are available at RNA-seq data are publicly available at the Gene Expression Omnibus (GEO) under accession number GSE355023.

## Supplemental Figures

**Fig. S1.**
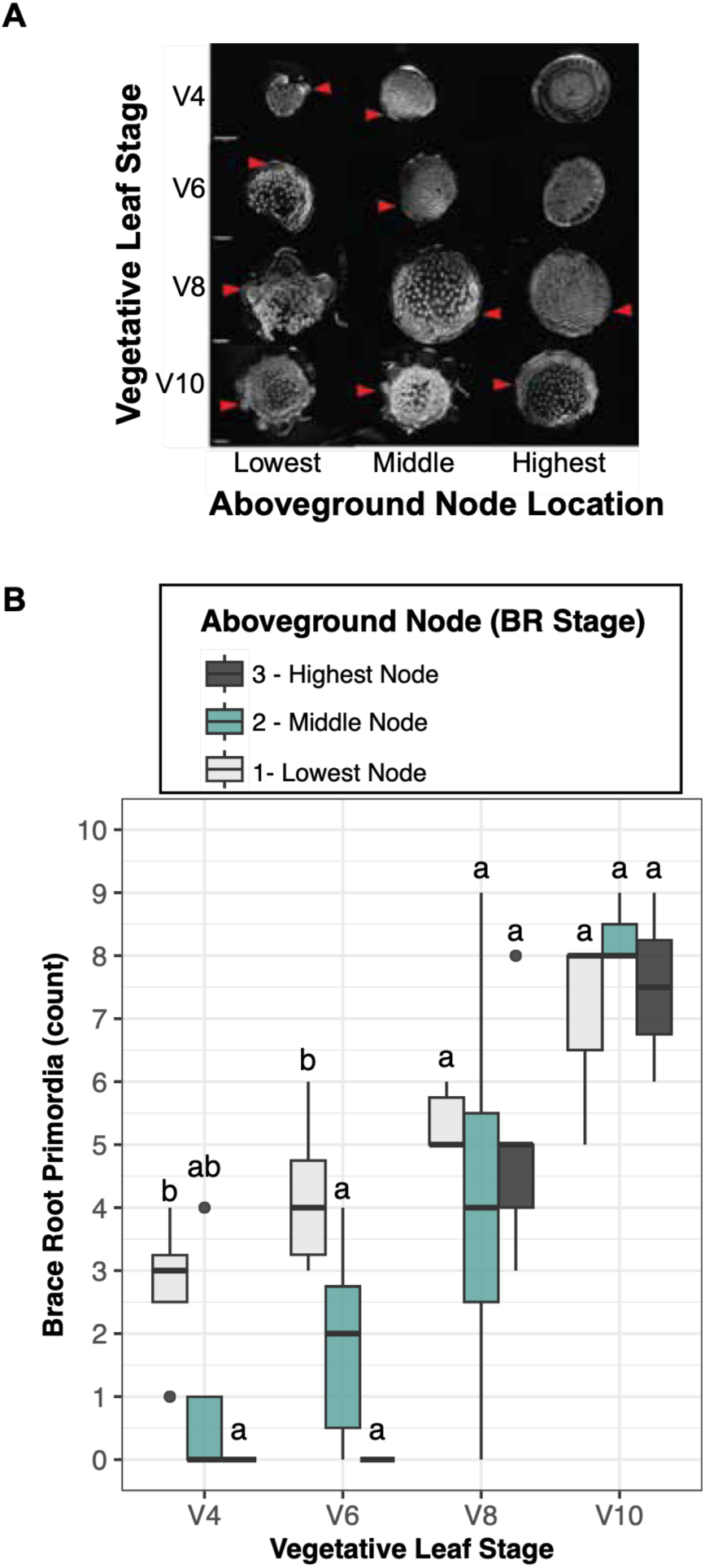
The spatiotemporal progression of brace root development. A) Analysis of brace root anatomy in B73 plants at the vegetative leaf 4 (V4), V6, V8, and V10 stages at the three aboveground nodes closest to the soil. Sections were stained with fuchsin and imaged using a Zeiss Axio Zoom microscope. Red arrowheads indicate brace root primordia or emerged brace roots. B) Quantification of brace root primordia shows two temporal trajectories: within individual nodes over time and sequentially across nodes within a growth stage. Data that share a letter are not statistically different from one another (*P* > 0.05).

**Fig S2.**
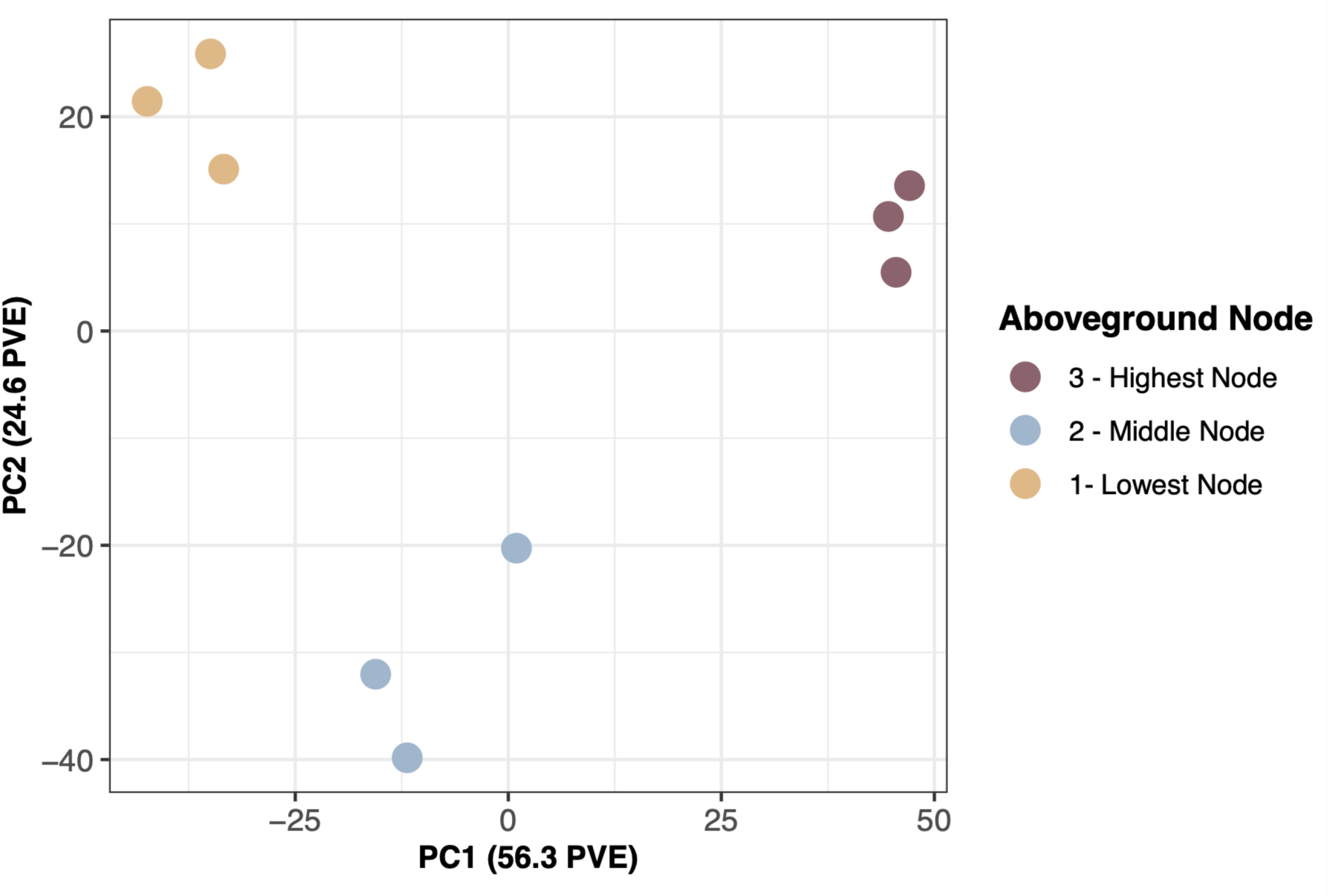
Brace root development has distinct transcriptional signatures. A principal component analysis (PCA) of transcriptome data from three V6 B73 plants shows that within nodes samples cluster together.

**Fig S3.**
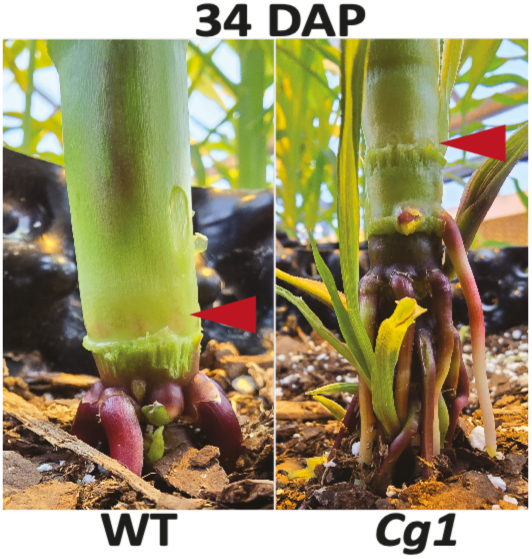
*Corngrass1 (Cg1)* develops one tier of brace roots at each node. Brace root primordia were characterized in the maize *Cg1* mutant and its wild-type sibling at 34 dap. While the *Cg1* develops brace roots at all nodes, it only develops one tier of brace roots. The red arrowheads indicate the pulvinus containing brace root primordia.

**Fig S4.**
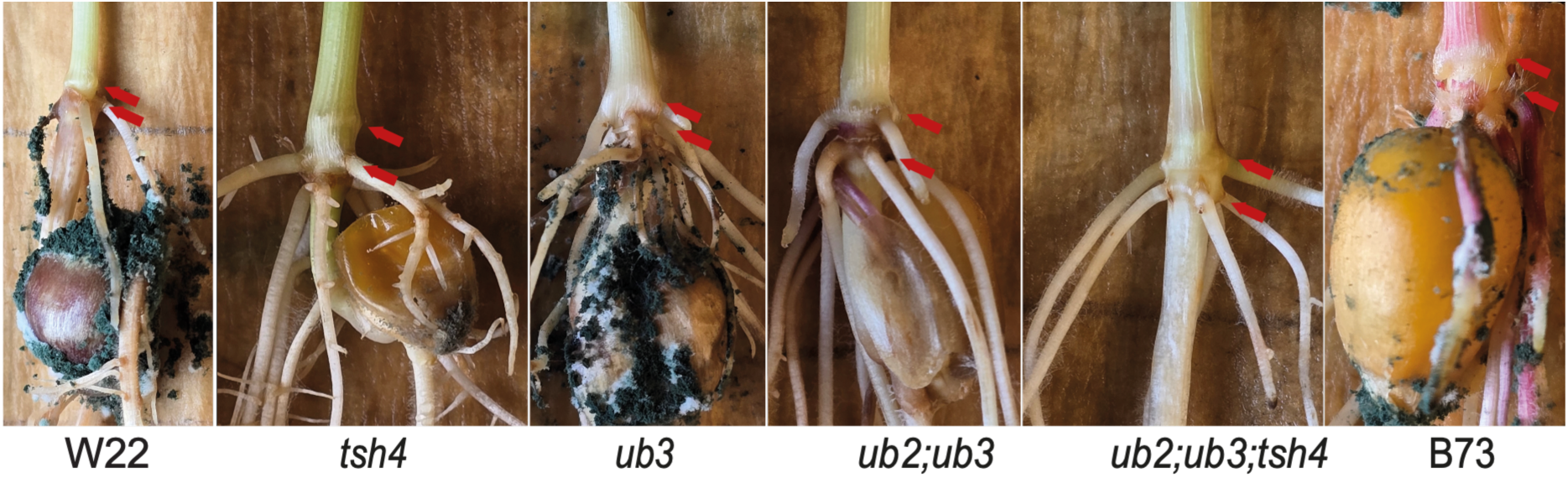
One tier of roots emerged from crown root nodes of *sbp* mutant plants. To determine if belowground nodal roots (i.e., crown roots) developed two tiers of roots at each node, images of plants taken at 11 dap were assessed. Unlike aboveground brace root nodes, only a single tier was observed for crown roots of *sbp* mutants. The red arrowheads indicate the coleoptile and crown root node.

**Fig S5.**
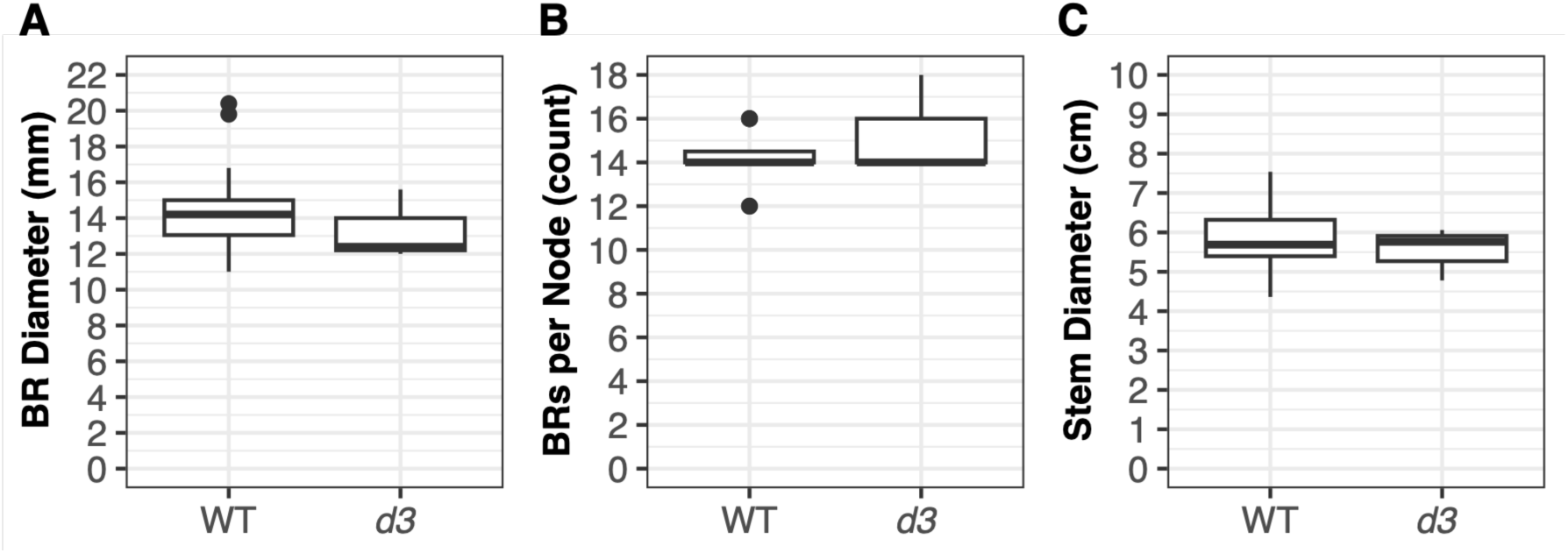
Brace root architecture does not differ in the *dwarf3* (*d3*) mutant. A) The size of emerged brace roots did not differ between *d3* and its wild-type sibling. B) There number of brace roots per node in *d3* compared to its wild-type sibling did not differ. C) The *d3* mutant and its wild-type sibling have similar stem diameters. Black dots indicate outliers. Statistical significance assessed at *P* < 0.05.

**Fig S6.**
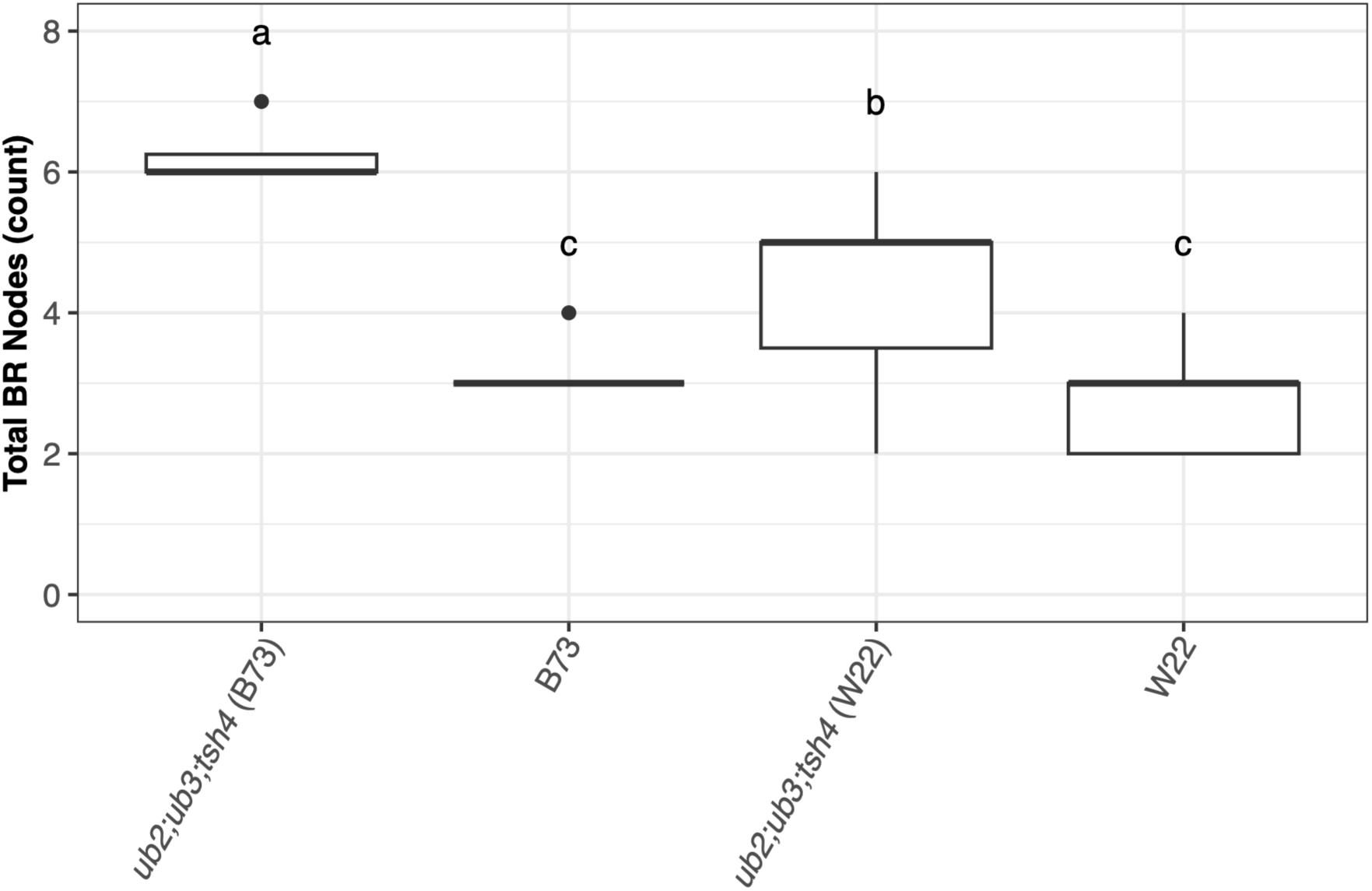
Inbred background affects the penetrance of *sbp* mutant phenotypes. The *tsh4;ub2;ub3* mutant in the B73 background has more nodes that produced brace roots than the same mutant in the W22 background. Both wild-type lines had similar numbers of nodes that produced brace roots. Black dots indicate outliers. Genotypes that share a letter did not significantly differ from one another when a post hoc Tukey honest significant difference (HSD) test was employed (*P*<0.05).

**Fig S7.**
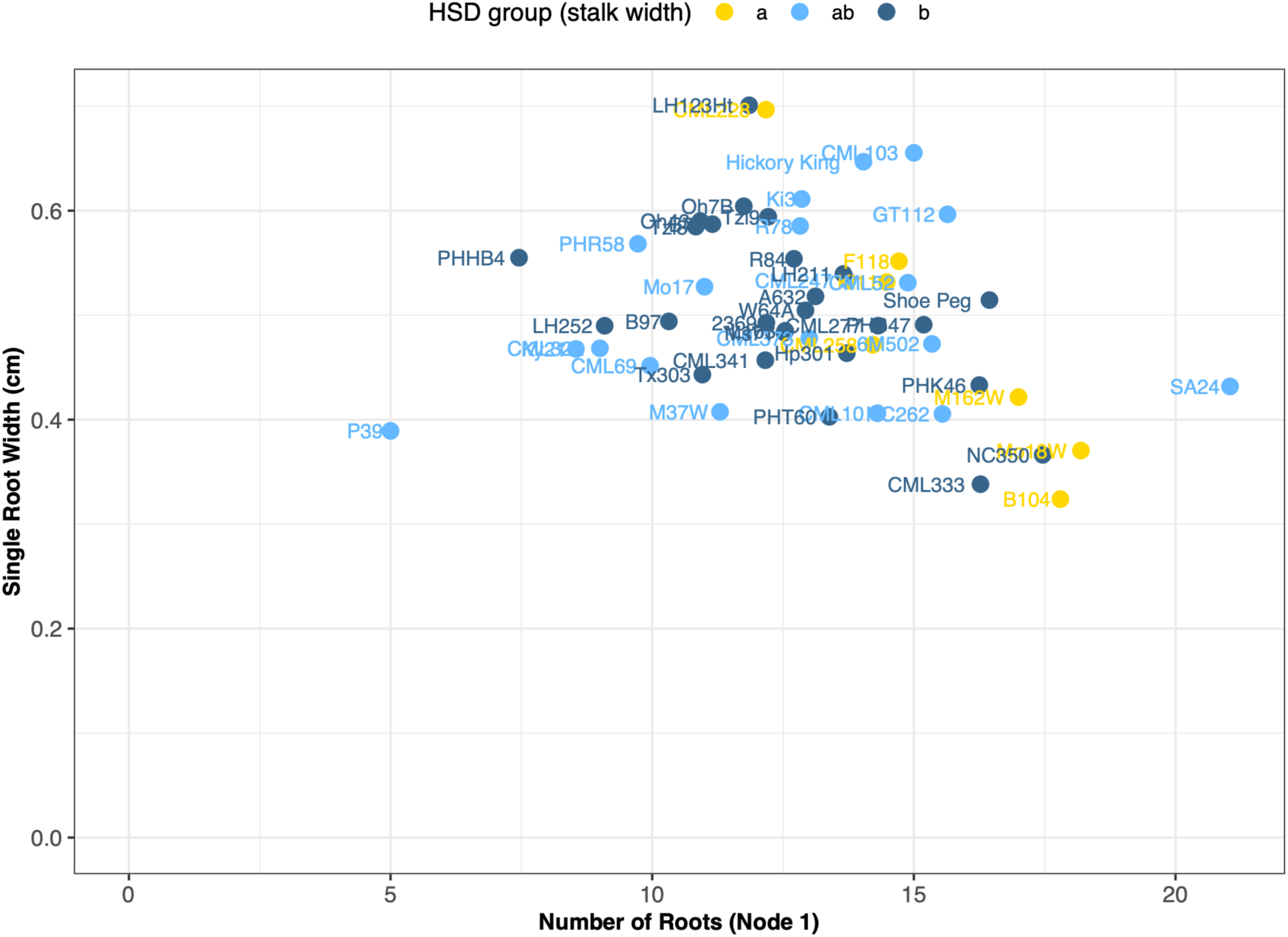
Inbred lines vary for the size and number of brace roots. Data from a previous study (Hostetler et al. 2022) was used to visualize the relationship between brace root size, brace root number, and stem size. Genotypes with the same-colored dot did not significantly differ for stem size when a post hoc Tukey honest significant difference (HSD) test was employed (*P*<0.05).

