## Supplementary material for "SBP transcription factors repress maize brace root initiation and regulate primordia expansion": Fig S1

A

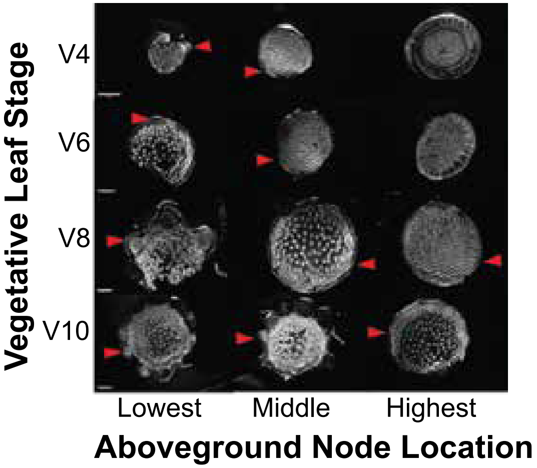

Vegetative Leaf Stage
V4
V6
V8
V10
Lowest
Middle
Highest
Aboveground Node Location


B

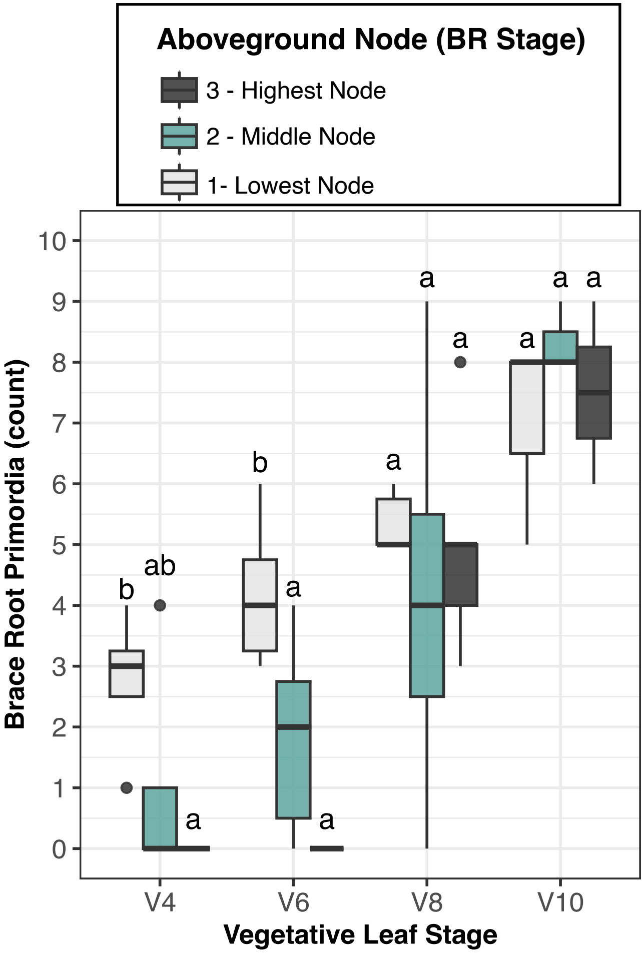

Aboveground Node (BR Stage)
3 - Highest Node
2 - Middle Node
1- Lowest Node
Brace Root Primordia (count)
0
1
2
3
4
5
6
7
8
9
10
V4
V6
V8
V10
Vegetative Leaf Stage
b
ab
a
b
a
a
a
a
a
a
a
a
