## Supplementary figures and images for "SBP transcription factors repress maize brace root initiation and regulate primordia expansion"

### Fig S2

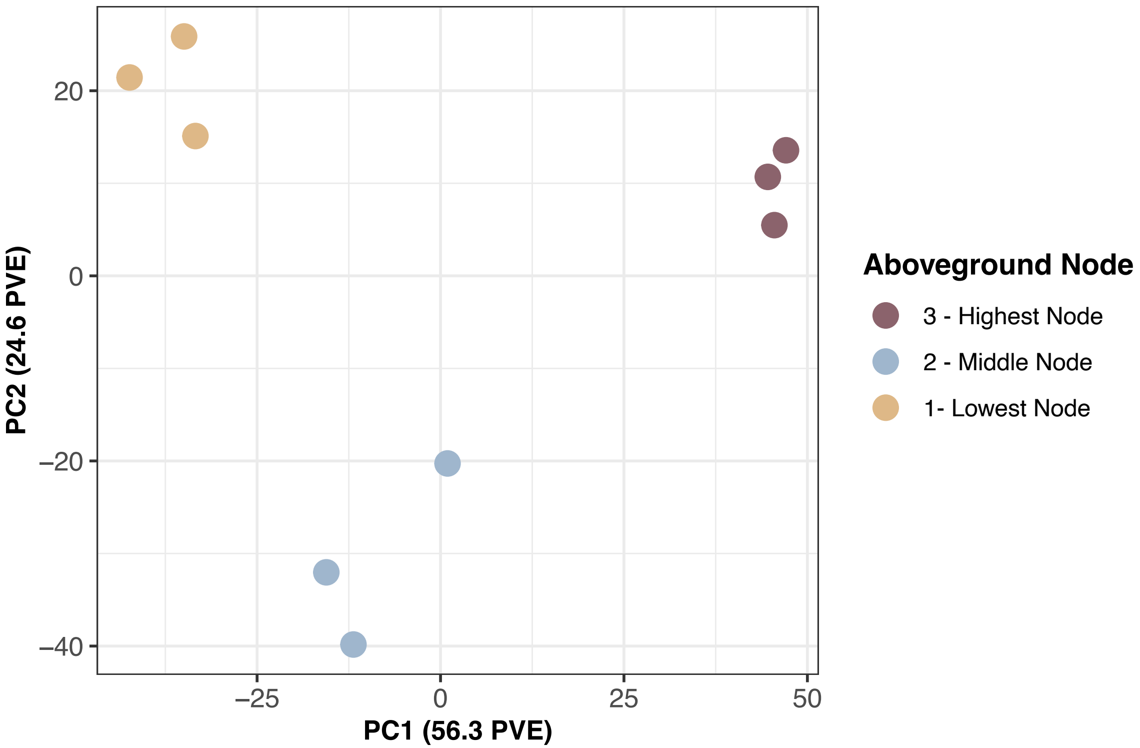
PC2 (24.6 PVE)
20
0
−20
−40
−25
0
25
50
PC1 (56.3 PVE)
Aboveground Node
3 - Highest Node
2 - Middle Node
1- Lowest Node

### Fig S3

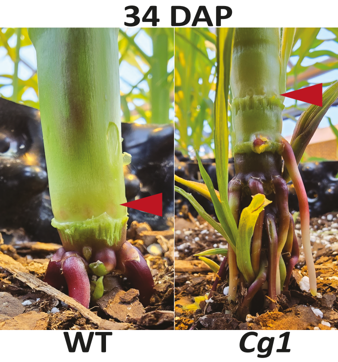
34 DAP
WT
Cg1

### Fig S4

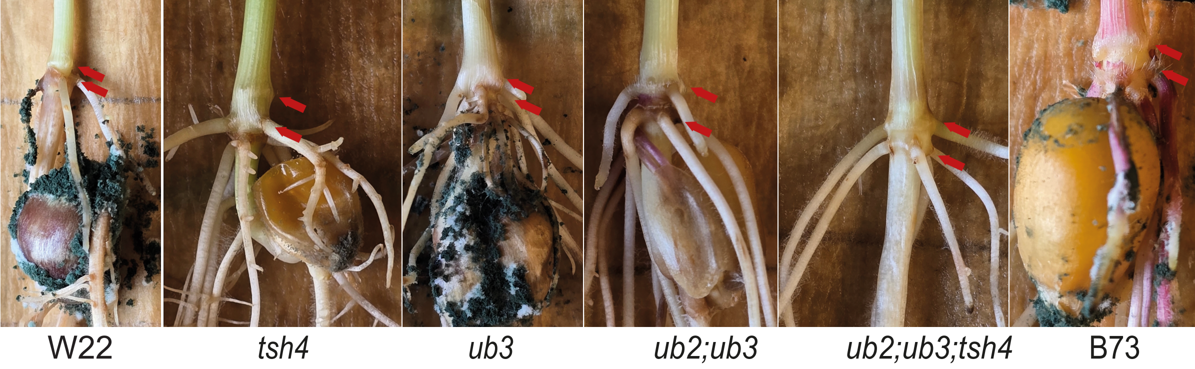
W22
*tsh4*
*ub3*
*ub2;ub3*
*ub2;ub3;tsh4*
B73

### Fig S5

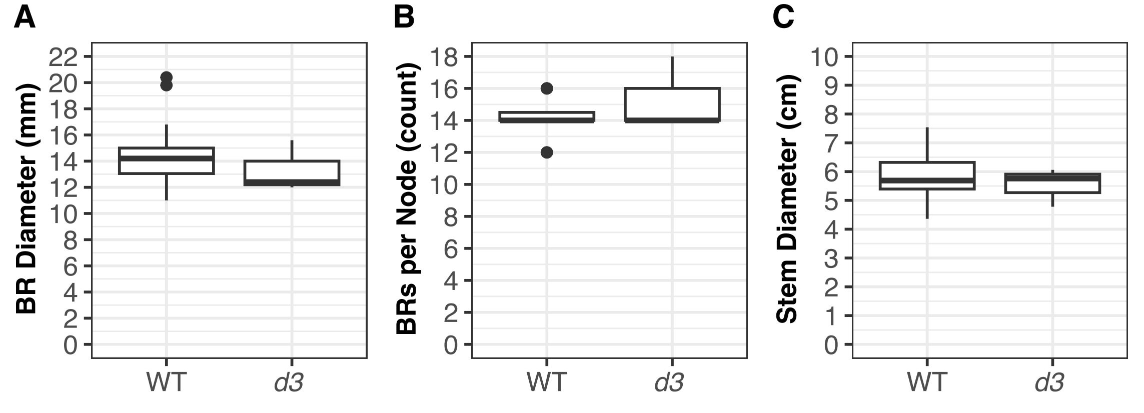
A
BR Diameter (mm)
0
2
4
6
8
10
12
14
16
18
20
22
WT
d3
B
BRs per Node (count)
0
2
4
6
8
10
12
14
16
18
WT
d3
C
Stem Diameter (cm)
0
1
2
3
4
5
6
7
8
9
10
WT
d3

### Fig S6

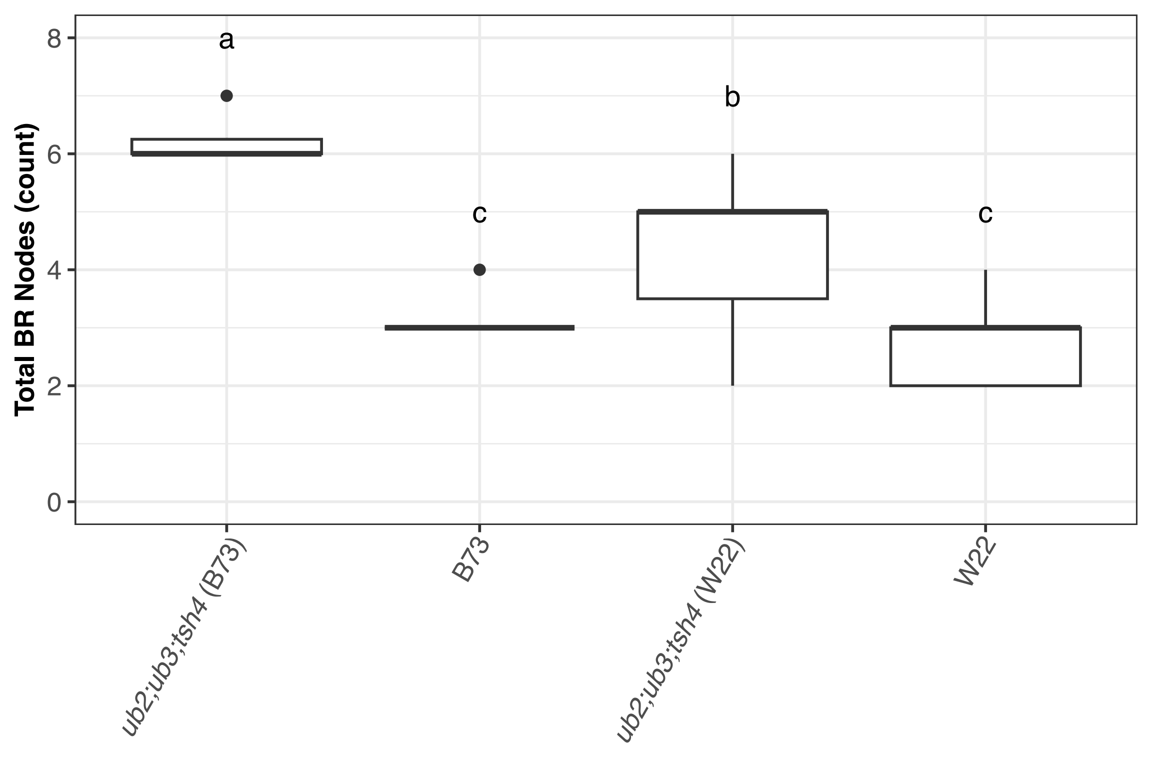

Total BR Nodes (count)
0
2
4
6
8
a
c
b
c
ub2;ub3;tsh4 (B73)
B73
ub2;ub3;tsh4 (W22)
W22
