## Supplementary material for "SBP transcription factors repress maize brace root initiation and regulate primordia expansion": Fig S7

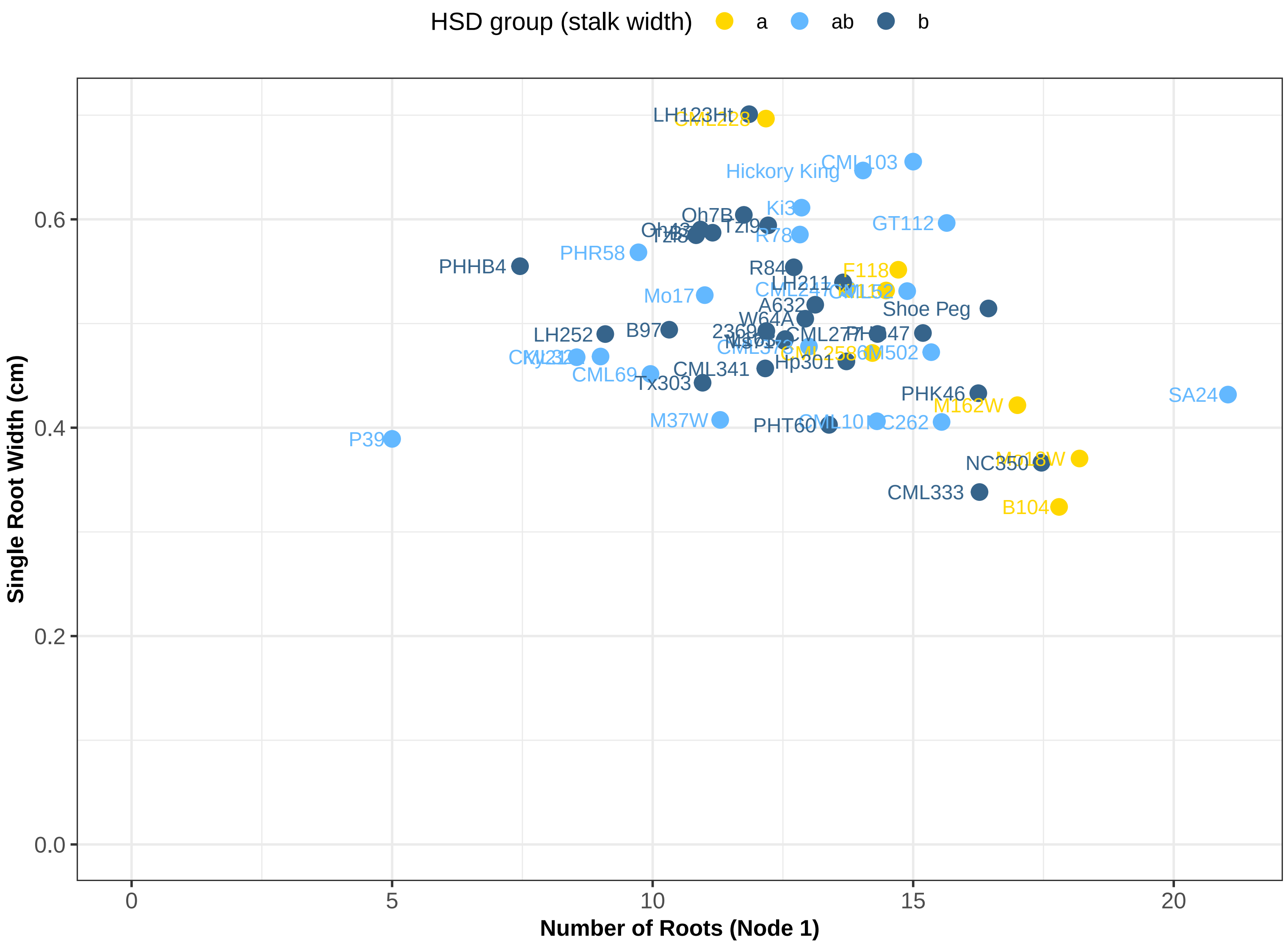

HSD group (stalk width)
a
ab
b
Single Root Width (cm)
Number of Roots (Node 1)
0.0
0.2
0.4
0.6
0
5
10
15
20
LH123Ht
CML103
Hickory King
Ki3
Oh7B
GT112
R78
PHR58
PHHB4
R84
F118
LH211
Mo17
A632
Shoe Peg
W64A
LH252
B97
M502
CML341
Hp301
CML69
Tx303
PHK46
SA24
M162W
M37W
PHT60
P39
CML333
B104
